# Ambivalent partnership of the Drosophila posterior class Hox protein Abdominal-B with the Extradenticle and Homothorax cofactors

**DOI:** 10.1101/2024.07.03.601496

**Authors:** Jesús R. Curt, Paloma Martín, David Foronda, Bruno Hudry, Ramakrishnan Kannan, Srividya Shetty, Samir Merabet, Andrew Saurin, Yacine Graba, Ernesto Sanchez Herrero

## Abstract

Hox proteins, a sub-group of the homeodomain (HD) transcription factor family, provide positional information for axial patterning in development and evolution. Hox protein functional specificity is reached, at least in part, through Pbc (Extradenticle (Exd) in Drosophila) and Meis/Prep (Homothorax (Hth) in Drosophila) cofactor interactions. Most of our current knowledge of Hox protein specificity stems from the study of anterior and central Hox proteins, identifying the molecular and structural bases for Hox/Pbc/Meis-Prep cooperative action. Posterior Hox class proteins, Abd-B in Drosophila and Hox9-13 in vertebrates, have been comparatively less studied. They strongly diverge from anterior and central class Hox proteins, with a low degree of HD sequence conservation and the absence of a core canonical Pbc interaction motif. Here we explore how Abd-B function interface with that of Exd/Hth using several developmental contexts, studying mutual expression control, functional dependency and intrinsic protein requirements. Results identify cross regulatory interactions setting relative expression and activity levels required for proper development. They also reveal organ-specific requirement and a binary functional interplay with Exd and Hth, either synergistic or antagonistic. This highlights context specific use of Exd/Hth cofactors, and a similar context specific use of Abd-B protein intrinsic protein requirements.

## INTRODUCTION

Gene regulation is central for implementing differences between cells of an organism sharing the same genome during development, evolution and when switches to pathological situations occur. Gene regulation is largely mediated by transcription factors that bind promoters or more distant gene regulatory regions to activate or repress gene expression, the first step towards imprinting differences in protein content within cells. A large number of transcription factors are mobilized for proper gene regulation, in the range of 630 in *Drosophila* to 1500 human. The large number of transcription factors contrasts with the limited number of strategies evolved to bind DNA, resulting in the grouping of transcription factors in only a few families (Lambert et al., Cell 2018). The most prominent of these is the zinc finger family, representing almost half of all transcription factors. In this specific instance, large variations in the type and number of zinc finger allow generating a large repertoire of binding specificity. The second most prominent family of transcription factors is the homeodomain (HD) transcription factor family, with >200 representatives in humans. The HD is a helix-turn-helix DNA binding domain usually 60 amino acid long and highly conserved, in particular within the DNA recognition helix that make strong contacts with DNA (Gehring et al., 1994). The high degree of HD conservation endows HD transcription factors with little DNA binding specificity, contrasting with their diverse and specific biological functions (Bobola and Merabet, 2017).

Hox proteins, a sub-group of the HD transcription factor family, have provided a paradigm over years to address the discrepancy between similar DNA binding and specificity in function (Noyes et al., 2008; Berger et al., 2008; Mann et al., 2009). Hox genes are axially differentially expressed along the antero-posterior axis, providing positional information for axial patterning in development and evolution (Lewis, 1978; Rezsohazy et al., 2015). A solution to the specificity paradox comes from the observation that Hox proteins acts along with cofactors or partner proteins, and that the resulting complexes display higher DNA binding specificity. The best studied cofactors, Pbc and Meis/Prep proteins, are also HD containing proteins, and form dimeric/trimeric DNA binding complexes with increased DNA binding specificity (Jacobs et al., 1999; Ryoo et al., 1999; Shen et al., 1999; Shanmugan et al, 1999; Ferretti et al., 2000; Vlachakis et al., 2001; Mann et al., 2009), revealing “latent” specificity of Hox proteins (Joshi et al., 2007; Slattery et al., 2011, Merabet and Mann, 2016). The interactions between Pbc and Hox proteins have been studied by biochemical (Chang et al., 1995; Johnson et al., 1995; Phelan et al., 1995) and structural (Passner et al, 1999; Piper et al., 1999; LaRonde –Leblanc and Wolberger, 2003; Foos et al., 2019) methods, and studies in mouse and Drosophila have shown that the presence of Exd/Pbx cofactors helps to discriminate targets Hox proteins regulate in vivo (Sánchez-Higueras et al., 2019; Bridoux et al., 2020; Merabet and Mann, 2016).

In Drosophila, Pbc and Meis/Prep class proteins have a single representative, Extradenticle (Exd) and Homothorax (Hth) respectively, making the genetic analysis easier than in vertebrates (four Pbx, three Meis and two Prep proteins in mouse; Bobola and Sagerström, 2024). Mutations in *exd* and *hth* modify Hox gene activity without changing Hox expression (Peifer and Wieschaus, 1990; Rauskolb et al., 1993; Rieckhof et al., 1997; Kurant et al., 1998; Pai et al., 1998). Early studies established that the formation of a Hox dimeric/trimeric complex was mediated by a short protein motif N-terminal to the HD known as the Hexapeptide (HX) (Mann and Chan, 1996; Mann et al., 2009). Subsequent studies demonstrated this motif was dispensable for some Pbc-dependent Hox functions (Galant et al., 2002; Merabet et al., 2003) and that the region immediately C-terminal to the HD also contacts Exd (Merabet et al., 2009; Lelli et al., 2011; Saadaoui et al., 2011; Hudry et al., 2012; Foos et al., 2015: Dard et al., 2018; Dard et al., 2019; Merabet and Hudry, 2011; Merabet and Mann, 2016; Bobola and Merabet, 2016; Ortíz-Lombardía et al., 2017; Singh et al., 2020).

Based on the homeodomain protein sequence and expression along the antero-posterior axis, Hox paralogs have been classified as belonging to the anterior, central or posterior groups. In vertebrates, the posterior group comprises Hox genes 9-13 while in *Drosophila* the group includes just one gene, *Abdominal-B* (*Abd-B*). Hox9-10 proteins have a rudimentary HX whose sequence diverges for a core consensual HX motif, yet it displays a central W), and Hox11-13 lack the HX (Duboule, 1994)The lack (or strong degeneration) of the initially identified Pbc interacting motif likely limited the study of how Pbc class (and Meis/Prep classes) functionally interface with posterior class Hox proteins. Most of our current knowledge pertaining to mechanisms of Hox protein specificity thus stems from the study of anterior and central Hox proteins, identifying modes of functional interactions as well as the molecular and structural bases for cooperative action.

*Abd-B* specifies the posterior abdominal segments and some terminal structures in embryo and the posterior abdomen and the genitalia in adults (Sanchez-Herrero et al. 1985; Tiong et al., 1985; Casanova et al., 1986). In the *Drosophila* embryo, *Abd-B* represses *exd* and *hth* expression, so that in the last abdominal segment (A8-A9) levels of these two proteins are very low (Rauskolb et al., 1993; Rieckhof et al., 1997; Azpiazu and Morata, 1998; Kurant et al., 1998; Rivas et al., 2013; Sambrani et al., 2013). If *exd* and *hth* expression is maintained in the posterior of the embryo, Abd-B function is impaired, binding of the Hox protein to characterized enhancers is disturbed, and an aberrant phenotype in posterior spiracles is observed (Rivas et al., 2013; Sambrani et al., 2013). These genetic and molecular studies argue for a functional antagonism between the posterior Hox protein Abd-B and Exd/Hth, indicative of a relationship distinct from the one described for anterior and central Hox proteins. However, requirement of *exd/hth* for *Abd-B* to make ectopic embryonic abdominal denticles (Sambrani et al., 2014) suggests a more complex interplay. Here we explore how Abd-B function interfaces with that of Exd/Hth using several developmental contexts, studying mutual expression control, functional dependency and intrinsic protein requirements.

## RESULTS

### Lack of Abd-B-mediated repression results in Abd-B, Hth and Exd co-expression in the A7 pupal abdomen

Adult Drosophila males have only six abdominal segments as a result of abdominal segment 7 (A7) suppression by Abd-B in pupae (Sánchez-Herrero et al. 1985; Tiong et al., 1985). In the wild type the A7 is extruded at about 35-42 hours after puparium formation (APF) while in *Abd-B* mutants an A7 is observed (Yoder et al., 2011, Foronda et al., 2012). As previously described (Kopp and Duncan, 2002; Wang et al., 2011; Foronda et al., 2012; Singh and Mishra, 2014), there are higher levels of *Abd-B* in A7 than in A6 pupal segments (Fig. 1A). Exd and Hth are expressed in all the nuclei of the pupal abdomen including Larval Epidermal Cells (LECs) and histoblasts, and contrary to what happens in the embryo, where Abd-B down-regulates *exd* and *hth* expression (Azpiazu and Morata 1998; Rivas et al., 2013; Sambrani et al., 2013), there is co-expression of Abd-B with Exd and Hth, even in A7, where Abd-B levels are higher (Fig. 1B; Supplementary Fig. 1). This defines a biological context where Abd-B, Exd and Hth are co-expressed, allowing to study how they functionally interface.

**Figure 1.**
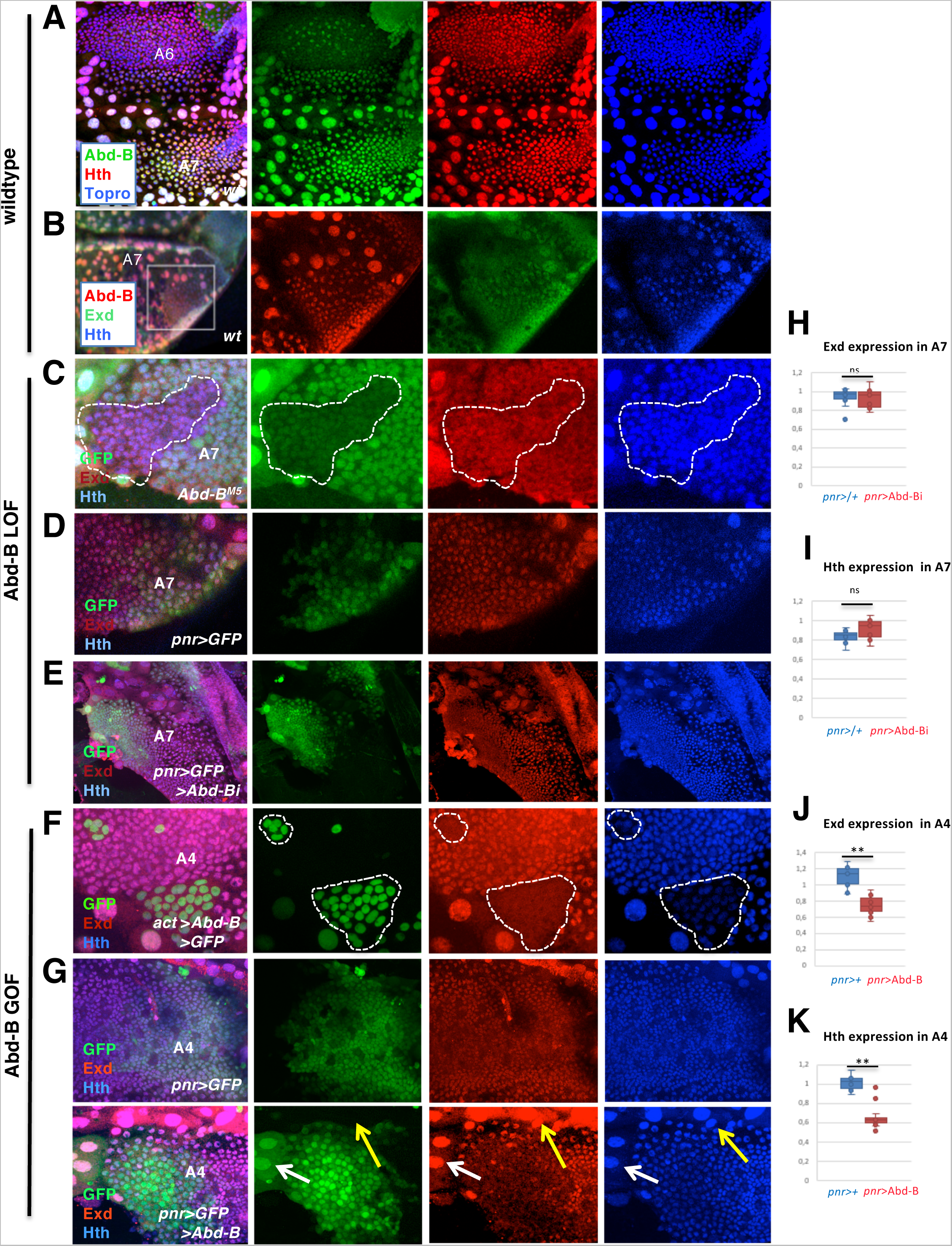
*Abd-B, exd* and *hth* wildtype expression and regulation of *exd* and *hth* by *Abd-B.* **(A)** Male wildtype pupa showing higher levels of Abd-B expression (in green) in the A7 than in the A6. Hth is in red and Topro is in blue**. (B)** Male A7 segment of a pupa showing the coincident expression of Abd-B (red), Exd (green) and Hth (blue) in histoblasts and Larval Epidermal Cells). **(C)** *Abd-B^M5^*clones, induced in the pupal male A7 and marked by the absence of GFP, showing there is no major change in Exd (red) or Hth (blue) expression with respect to wildtype cells. **(D)** In *pnr*-Gal4 UAS-GFP males, the *pnr* domain of the A7 segment has similar levels of *hth* to those of the *pnr^-^* domain of the same segment. **(E)** The reduction of *Abd-B* expression in the *pnr* region of the male A7, marked with GFP (*pnr*-Gal4 UAS-GFP UAS-Abd-BRNAi), does not change Hth levels of expression with respect to the domain not reducing Abd-B expression. **(F)** The ectopic expression of Abd-B (line 1.1) in the pupal A4 segment, marked by the GFP expression, reduces the expression of Exd (red) and Hth (blue) with respect to adjacent, wildtype cells. **(G)** In *pnr*-Gal4 UAS-GFP pupae, the A4 segment has similar levels of expression of Exd (red) and Hth (blue’) in the *pnr^+^* (marked with GFP’) and the *pnr*^-^ domains. In *pnr*-Gal4 UAS-GFP UAS-Abd-B (line 1.1), by contrast, the levels of expression of Exd (red) and Hth (blue’) in the *pnr^+^* domain, marked with GFP are substantially reduced in comparison with the *pnr*^-^ domain. The white arrow indicate LECs that express GFP (and therefore ectopically express *Abd-B*) but do not reduce Exd or Hth levels as compared with LECs that do not activate Abd-B (yellow arrow). **(H, I)** Quantification of the ratio of expression in a central (pnr+) with respect to a more lateral (pnr-) domain of Exd (H) or Hth (I) in *pnr*-Gal4/*+* and *pnr*-Gal4 UAS-Abd-BRNAi histoblast nests of the A7 segments of male pupal abdomens. **(I)** Quantification of the ratio of expression in a central (pnr+) with respect to a more lateral (pnr-) domain of Hth an *pnr*-Gal4/*+* and *pnr*-Gal4 UAS-Abd-BRNAi histoblast nests of the A7 segments of male pupal abdomens. **(J, K)** Quantification of the ratio of expression in a central (pnr+) with respect to a more lateral (pnr-) domain of Exd (J) or Hth (K) in *pnr*-Gal4/*+* and *pnr*-Gal4 UAS-Abd-B histoblast nests of the A3-A4 segments of male pupal abdomens. Pupae in all panels are of about 24-30h APF. Statistical analysis of the data in H-K was done by two-tailed t-tests, with n=10-12 pupae.

To analyze the interaction between *Abd-B* and *exd*/*hth*, we manipulated the expression levels of either *Abd-B* or *exd*/*hth*, and observed the expression of the other. If *Abd-B* expression is reduced in the male A7 by inducing *Abd-B* mutant (*Abd-B^M5^*) clones, *hth* and *exd* expression does not significantly change (Fig. 1C). Analyzed at about 24-30h APF, we found that 36/44 of these clones showed similar Hth or Exd levels to those of cells outside the clones (Fig. 1C). However, the remaining 8 clones showed a weak increase of expression (Supplementary Fig. 2), highlighting variability in the response to AbdB loss, which could reflect distinct timing of clonal loss of Abd-B, which in turn may also affect the level of AbdB protein reduction.

We also used local *Abd-B* inactivation, combining the GAL4/UAS system with the *pannier* (*pnr*)-Gal4 line, active in the central dorsal abdominal region (Calleja et al., 2000), and the UAS-Abd-B RNAi, allowing to compare expression levels in the central (*pnr+*) and lateral (*pnr*-) regions of the same segment and score the *pnr*+/*pnr*-gene levels ratio as a measure of gene regulation, both in experimental and control (*pnr*-Gal4/+) animals. Conditional *pnr-*Gal4 driven expression was achieved using the expression of a thermosensitive form of Gal80 (Gal80^ts^). In our experimental conditions (see Methods), expression of Abd-B RNAi does not impact, at about 28-32h APF, on Exd and Hth expression in histoblasts in the A7 segment (Fig. 1E, compare with D, H, I). At later stages (about 34-38h APF), an increase in *hth* expression is seen in pupae where *Abd-B* is reduced in the A7 segment with the *Abd-B^MD761^* Gal4 line, a Gal4 line driving expression in the A7 (see Methods and below), but such increase is also observed in *Abd-B^MD761^* UAS-*GFP*/+ control pupae (Supplementary Fig. 3).

The forced expression of *Abd-B* in the anterior abdominal segments however, reduces *hth* and *exd* expression: in *Abd-B*-expressing clones in the A3-A5 segments of about 24-28h APF pupae Exd and Hth expression is strongly reduced in histoblasts while expression in large epidermal cells (LECs) does not seem affected (Fig. 1F). Similarly, in *pnr*-Gal4 UAS-GFP *tub*-Gal80^ts^ UAS-Abd-B pupae a reduction of Hth and Exd expression in the *pnr* domain of A3-A5 segments is observed in the histoblasts (but not in LECs), as compared to controls (Fig. 1G, J, K). Thus, while wild type expression of *Abd-B* in A7 does not repress Exd and Hth, its ectopic expression in A3-A5 does.

Collectively, the expression and regulatory interactions studies show there is co-expression of the three proteins in the pupal male A7 segment while in the embryo the repression of *exd* and *hth* by Abd-B prevents the co–expression of the three proteins (Azpiazu and Morata, 1998; Rivas et al., 2013; Sambrani et al., 2013). This difference results from lack of Abd-B-mediated repression on *exd* and *hth* expression in the pupal abdomen, which seems dependent on a precise AbdB expression levels and/or on different Abd-B regulatory potential at distinct time and location

### Exd/Hth positively and contextually impacts *Abd-B* expression

Then, we studied if changes in Exd or Hth could affect Abd-B expression. When we induced and analyzed in the A7 clones mutant for *hth^P2^*, a null or close to null allele of *hth*, we observed a reduction of *Abd-B* expression in about half of the male A7 clones (8/18) (Fig. 2A). Similarly, in flip-out clones expressing an *hth RNAi* construct we also found 8/25 clones in the male A7 with reduced *Abd-B* expression (Fig. 2B). The clones that reduce *Abd-B* expression are predominantly located in the more posterior region of the segment.

**Figure 2.**
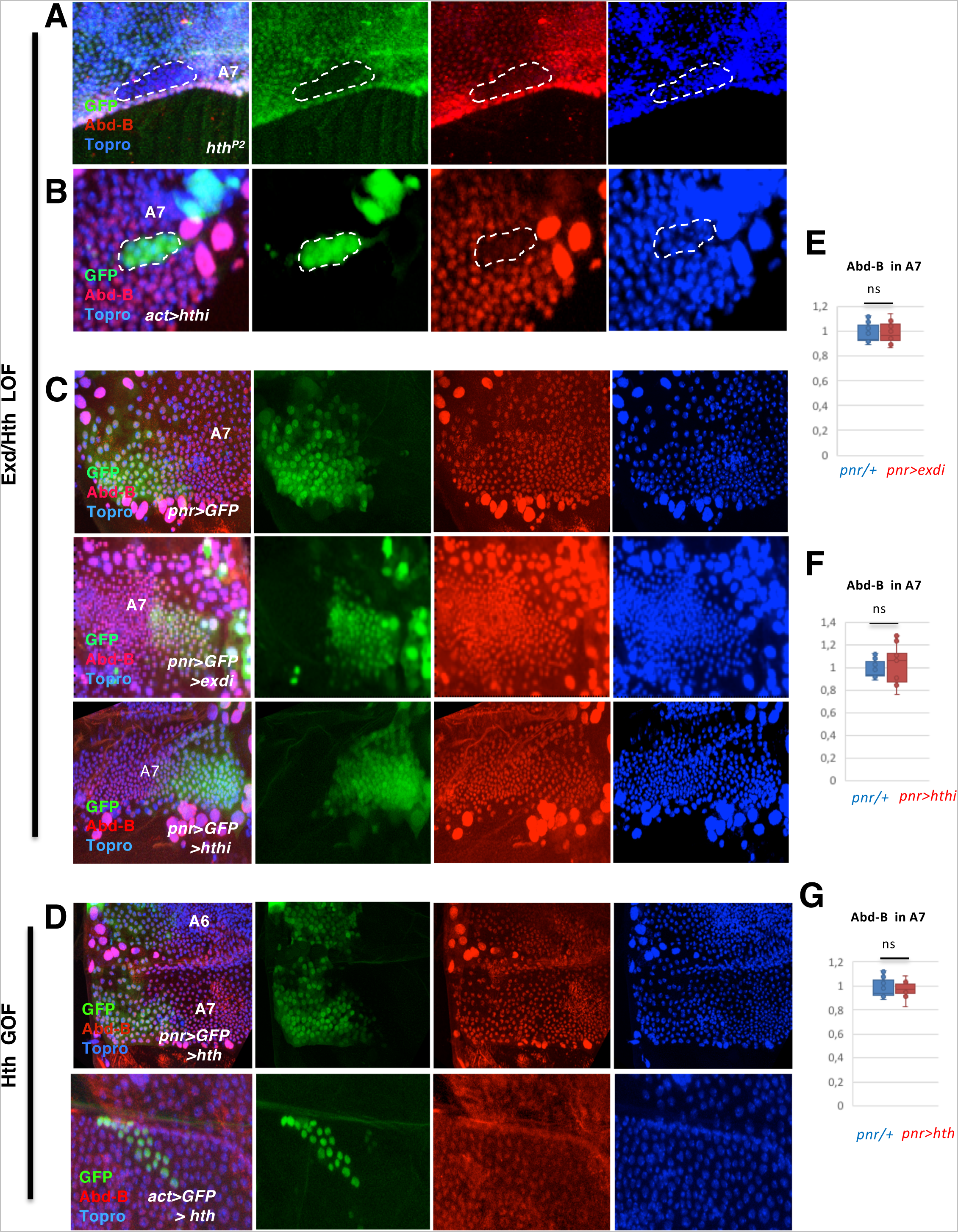
Regulation of *Abd-B* by *exd* and *hth* in the male pupal abdomen. **(A)** *hth^P2^*mutant clone in the pupal A7, marked by the absence of GFP, showing reduced expression of Abd-B (red). Topro is in blue. **(B)** Flip-out clone marked with GFP and expressing an hthRNAi construct (act>stop>Gal4 UAS-GFP UAS-hthRNAi) in the male A7 segment showing reduced expression of Abd-B (red). Topro, marking nuclei, is in blue. **(C)** In the male A7 segment of *pnr*-Gal4 UAS-GFP pupae the cells in the *pnr* domain have similar levels of Abd-B (in red) as those not expressing *pnr*. Topro, marking nuclei, is in blue’. If either *exd* or *hth* expression is reduced in the *pnr* domain, marked with GFP, of the male A7 (*pnr*-Gal4 UAS-GFP UAS-exdRNAi or *pnr*-Gal4 UAS-GFP UAS-hthRNAi), the expression of Abd-B is not significantly altered with respect to cells in the domain that does not express *pnr*. **(D)** male A7 segment of a *pnr*-Gal4 UAS-*hth* pupa showing similar levels of Abd-B expression (red) in the *pnr+* and *pnr-* domains (upper panels). Flip-out clone, marked with GFP, induced in the male A7 segment and expressing *hth* (lower panels). The levels of Abd-B in the clone (in red) do not change with respect to surrounding cells. **(E-G)** Quantification of the ratio of expression in a central (*pnr*+) with respect to a more lateral (*pnr*-) domain of Abd-B in *pnr*-Gal4/*+ pnr*-Gal4 UAS-exdRNAi (E), *pnr*-Gal4 UAS-hthRNAi (F) and *pnr*-Gal4 UAS-*hth* (G) in histoblast nests of the A7 segments of male pupal abdomens. Pupae in all panels are of about 24-30h APF except in A, which was of about 34h APF. Statistical analysis of the data in E-G was done by two-tailed t-tests, with n=10-12 pupae.

We also analyzed Abd-B expression in the A7 central (*pnr*) and lateral domains of *pnr*-Gal4 UAS-GFP *tub*-Gal80^ts^ UAS-exd RNAi, or UAS-hth RNAi 28-32h APF pupae, and compared the *Abd-B* expression with similarly treated controls. 10 out of 11 pupae of the *pnr*-Gal4 UAS-exd RNAi genotype and 14/15 of the *pnr*-Gal4 UAS-hth RNAi genotype had similar Abd-B levels of expression in both domains (*pnr^+^*and *pnr^-^*) of this segment, similarly to what is observed in the *pnr*-Gal4/+ controls (Fig. 2C, E, F). The *pnr*-induced loss of Exd or Hth thus seems inefficient in affecting Abd-B expression, suggesting that this mode of down regulation does not sufficiently lower Exd and Hth protein levels. Next, we studied the impact on Abd-B expression of over expression of *hth* in the *pnr* domain. This forced expression does not change the Abd-B levels in A7 with respect to the adjacent, *pnr*-negative domain (Fig. 2D, G). Flip-out clones expressing *hth* in the A7 do not modify *Abd-B* expression either (Fig. 2D). The Hth gain-of-function experiments thus do not show any increase of Abd-B over its wild type levels in A7.

We thus concluded that Exd/Hth positively impact on Abd-B expression, a situation not seen in the embryo (Rieckhof et al., 1997), and only evidenced by the clonal loss of Exd/Hth. As observed for *exd/hth* regulation by *Abd-B*, the variability observed in *Abd-B* regulation by *exd/hth*, when it does, suggest the need for a precise Abd-B/Hth/Exd levels for proper developmental activity.

### Reduction or increase of *exd*/*hth* cause *Abd-B* loss of function phenotypes in the male A7

We studied next the role of *exd*/*hth* in the development of the posterior abdomen, the *Abd-B* domain. *Abd-B* functions in the posterior abdomen by imposing posterior abdominal identity and suppressing the emergence of an A7 segment in the adult male.

Previous analyses showed a transformation of posterior abdominal segments to a more anterior segment in clones or gynandromorphs mutant for *exd* (Rauskolb et al., 1995). We also found that reduction of *exd* through *pnr*-mediated RNAi gene inactivation in the central region of the abdomen, or through *hth* loss in *hth^P2^* mutant clones, results in the appearance of trichomes, a marker of anterior abdominal segments, in A6, suggesting a transformation of A6 towards a more anterior abdominal identity (Fig. 3B-C’, compare with A). However, an excess of *hth* in the A6 also causes anteriorwards transformation (Ryoo et al., 1999; Fig. 3D, D’). This shows that manipulating *exd*/*hth* levels in either direction results in an *Abd-B*-like phenotype, thus highlighting a complex relationship between *Abd-B* and *exd*/*hth*, and indicating that *exd*/*hth* expression levels have to be tightly regulated in the posterior abdomen.

**Figure 3.**
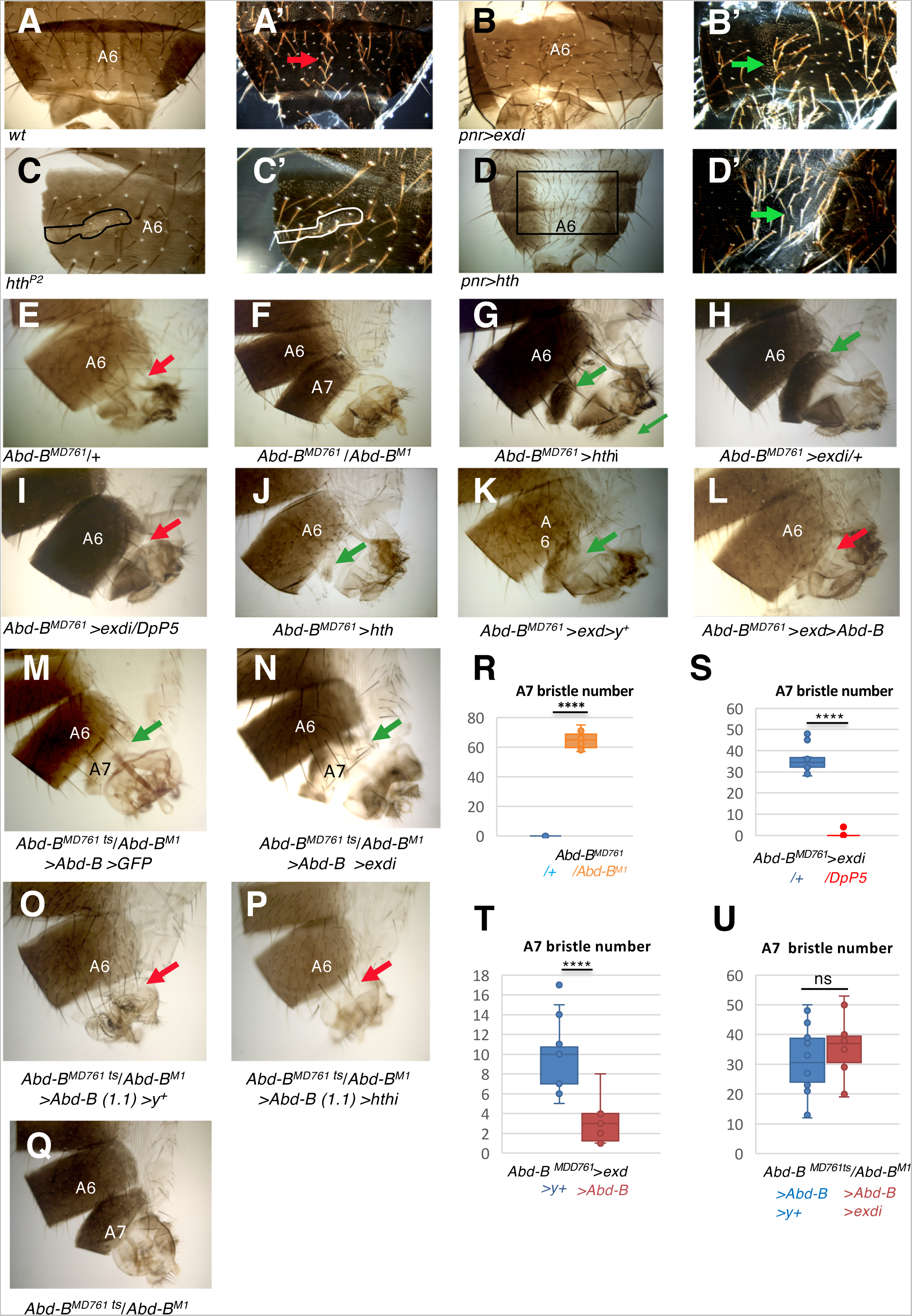
Reduction or excess of *hth*/*exd* cause anteriorwards transformations in the male abdomen. **(A, A’)** Bright (A) and dark field (A’) images of the dorsal A6 segment of a wildtype male, showing no trichomes in the medial region of the segment (red arrow). **(B, B’)** In *pnr*-Gal4 UAS-exdRNAi males, the dorsal central domain of the A6 segment, where *pnr* is expressed, presents many trichomes (B’, dark field image, green arrow), suggesting transformation to a more anterior segment. **(C, C’)** Bright (C) and dark field (C’) images of a clone mutant for *hth^P2^*, marked with *yellow* and outlined, showing trichomes within the clone, suggesting transformation to a more anterior segment. **(D, D’)** In the *pnr*-Gal4 UAS-*hth* abdomen, the central dorsal region, where *pnr* is expressed, present depigmentation and a great number of trichomes (inset of D in D’, dark field, green arrow), again suggesting transformation of the A6 to a more anterior segment. **(E)** *Abd-B^MD761^*/+ males have no A7 segment, like the wildtype. In this and subsequent panels the red arrows indicate absence of A7 and the green arrows presence of this segment. **(F)** *Abd-B^MD761^* /*Abd-B^M1^* male (mutant background for rescue experiments). The A7 is extensively transformed into the A6. (**G-H**) The reduction of either *hth* **(G)** or *exd* **(H)** results in the development of a small A7 segment. The formation of this segment depends on the amount of Abd-B since an extra dose of *Abd-B* (with DpP5) in an *MD761*-Gal4 UAS-exdRNAi background reverts the mutant phenotype to the wildtype (compare H with I). **(J-L)** The increase in the amount of *hth* **(J**) or *exd* **(K)** also forms an A7 segment, and this development is prevented if *Abd-B* is concomitantly expressed **(L). (M, N)** The expression of *Abd-B* in a *tub*-Gal80^ts^/UAS-*Abd-B*; *Abd-B^MD761^* UAS-GFP/*Abd-B^M1^* male (shift from 18°C to 29°C at third larval stage) partially reverts the phenotype of the *Abd-B* mutant background (**M**); this phenotype does not significantly change if there is reduction of *exd* (UAS-exdRNAi/UAS-*Abd-B*; *Abd-B^MD761^ tub*-Gal80^ts^ /*Abd-B^M1^* male with the same temperature shift) (**N**). **(O)** UAS-*y*^+^/UAS-*Abd-B (1.1*); *Abd-B^MD761^ tub*-Gal80^ts^ /*Abd-B^M1^*male (shift from 18°C to 29°C at third instar) showing complete suppression of A7 development. **(P)** The reduction of *hth* expression in UAS-*Abd-B* (*1.1*)/*+*; *Abd-B^MD761^ tub*-Gal80^ts^ /*Abd-B^M1^* UAS-hthRNAi males, with a similar temperature shift, shows also no A7 segment. (**Q**) *Abd-B^MD761^ tub*-Gal80^ts^/*Abd-B^M1^* male (the mutant background for experiments in O, P), also shifted from 18°C to 29°C in third instar larva, showing transformation of A7 into A6. **(R-U)** Quantification of the number of bristles in the following genotypes: *Abd-B^MD761^/+* (n=11) and *Abd-B^MD761^* /*Abd-B^M1^* (n=14) (**R**), *Abd-B^MD761^* UAS-exdRNAi and *Abd-B^MD761^* UAS-exdRNAi DpP5 (n=10 for both genotypes) (**S**), *Abd-B^MD761^* UAS-*exd* UAS-*y^+^* (n=14) and *Abd-B^MD761^* UAS-exd UAS-*Abd-B* (n=10) (**T**), and *tub*-Gal80^ts^/UAS-*Abd-B*; *Abd-B^MD761^* UAS-GFP/*Abd-B^M1^* (n=18) and UAS-exdRNAi/UAS-*Abd-B*; *Abd-B^MD761^ tub*-Gal80^ts^ /*Abd-B^M1^* males (n=14) (**U**). Statistical analysis of the data in R-U was done by two-tailed t-tests.

In the wildtype or in heterozygotes for *Abd-B^MD761^* there is no A7 (Fig. 3E). *Abd-B^MD761^* (named as *MD761*-Gal4 in Foronda et al., 2012) is a P-Gal4 insertion in the *iab-7* regulatory region of *Abd-B* that drives expression in the A7 segment (histoblasts and LECs) and is also mutant for the *iab-7* function of *Abd-B,* required for A7 suppression (Foronda et al., 2012). Loss of this suppression (emergence of an A7) is obtained when combining *Abd-B^MD761^* with the *Abd-B^M1^* mutation (Fig. 3F). The size of the A7 segment can be assessed by the number of bristles in the A7 segment, providing an accurate estimate of the segment size, and therefore of Abd-B activity (Fig. 3R). A reduction of *exd* or *hth* by driving *exd* or *hth* RNAi expression with the *Abd-B^MD761^* driver produces a segment in the A7 position that also bears anterior identity, phenocopying (although not as strongly) the lack of *Abd-B* (Fig. 3G, H). As observed in A6 (Fig. 3D, D’), increased expression of *exd*, and to a lesser extent of *hth*, also results in the appearance of A7 segments (Fig. 3J, K).

These results suggest a genetic interaction between *exd*/*hth* and *Abd-B* in male A7 development. To assess further this relationship, we explored the dependency of the *exd*/*hth*-induced phenotypes upon *Abd-B* expression levels. We observed that the A7 that develops when there is a reduction of *exd* is completely suppressed by increasing the doses of *Abd-B* from 2 to 3 with the introduction of DpP5, a duplication for the Bithorax Complex (compare Fig.3H with 3I; 3S). This is consistent with the *Abd-B* down-regulation observed in some *hth^P2^* mutant clones (Fig. 2A). While an excess of *exd* or *hth* does not reduce *Abd-B* expression (Fig. 2D), an *Abd-B* loss of function phenotype is also observed when *exd* expression is increased in the A7 (Fig. 3K), suggesting a reduction of Abd-B activity. Consistently, the concomitant expression of *Abd-B* and *exd* reduces the A7 formed by an excess of *exd* (compare Fig. 3K with 3L; 3T). These results suggest two different mechanisms by which excess or reduction in *exd*/*hth* levels regulate Abd-B function, one by controlling its level of expression and the other by modulating its activity. .

We also assessed further the relationship between the *Abd-B* and *exd*/*hth* phenotypes by exploring dependency of the *Abd-B*-induced phenotypes upon Exd levels. To this end, we compared the rescuing effect of *Abd-B* in the development of male A7 in the presence and in the absence of *exd*. As described above, the *Abd-B^MD761^*/*Abd-B^M1^* combination results in the development of an A7 in the male (Fig. 3F). The expression of *Abd-B* in this mutant background (taking advantage of *Abd-B^MD761^* being a Gal4 line) partially rescued this transformation (compare 3F and 3M; the Gal4/Gal80^ts^ system was used in these rescuing experiments, see Methods). If *exd* expression is simultaneously reduced in this “rescuing” background, A7 development is not significantly altered (Fig. 3M, N, U). This result in isolation would suggest that *Abd-B* does not need *exd* for A7 development. However, the findings reported in Fig. 3F-H argue for a common role of Abd-B and Exd/Hth in A7 suppression and identity. In addition, the finding that both the loss and gain of Exd and Hth results in similar A7 phenotypes, which underlines the importance of relative Abd-B and Exd/Hth levels, rather suggest that in the rescuing conditions used, we do not modify sufficiently the Abd-B to Exd/Hth ratio to impact on the remaining Abd-B A7 suppressive function. Supporting the importance of this ratio, if higher levels of *Abd-B* (obtained using the UAS-Abd-B line 1.1; Castelli-Gair et al., 1994) are induced in the *Abd-B^MD761^*/*Abd-B^M1^*background, the mutant phenotype is completely rescued, and the effect is independent of the presence of *hth* (Fig. 3O-Q).

The importance of the Abd-B to Exd/Hth ratio is further supported by the analysis of *wingless* (*wg*), an Abd-B target in the abdomen. The expression of *wg* in the male A7 is suppressed by *Abd-B* (Wang et al., 2011; Foronda et al., 2012; Fig. 4A). If *exd* levels are reduced in this segment, ectopic *wg* expression is also observed (Fig. 4A). We have found, however, Wg antibody expression in the male A7 sometimes difficult to observe precisely due to high background and to the curvature of the posterior abdomen, so we turned to analyze Abd-B activity by observing the expression of a Wg-GFP protein trap (Port et al., 2014) and in the A3-A4 abdominal segments. We found that by increasing *Abd-B* levels in the central dorsal abdomen with *pnr*-Gal4, Wg-GFP expression in the male A3-A4 segments is strongly reduced (Fig. 4B, C). Increasing Exd levels in this context impairs Abd-B ability to down-regulate Wg-GFP expression, suggesting high levels of Exd counteract the repressing activity of Abd-B (Fig. 4B, C).

**Figure 4.**
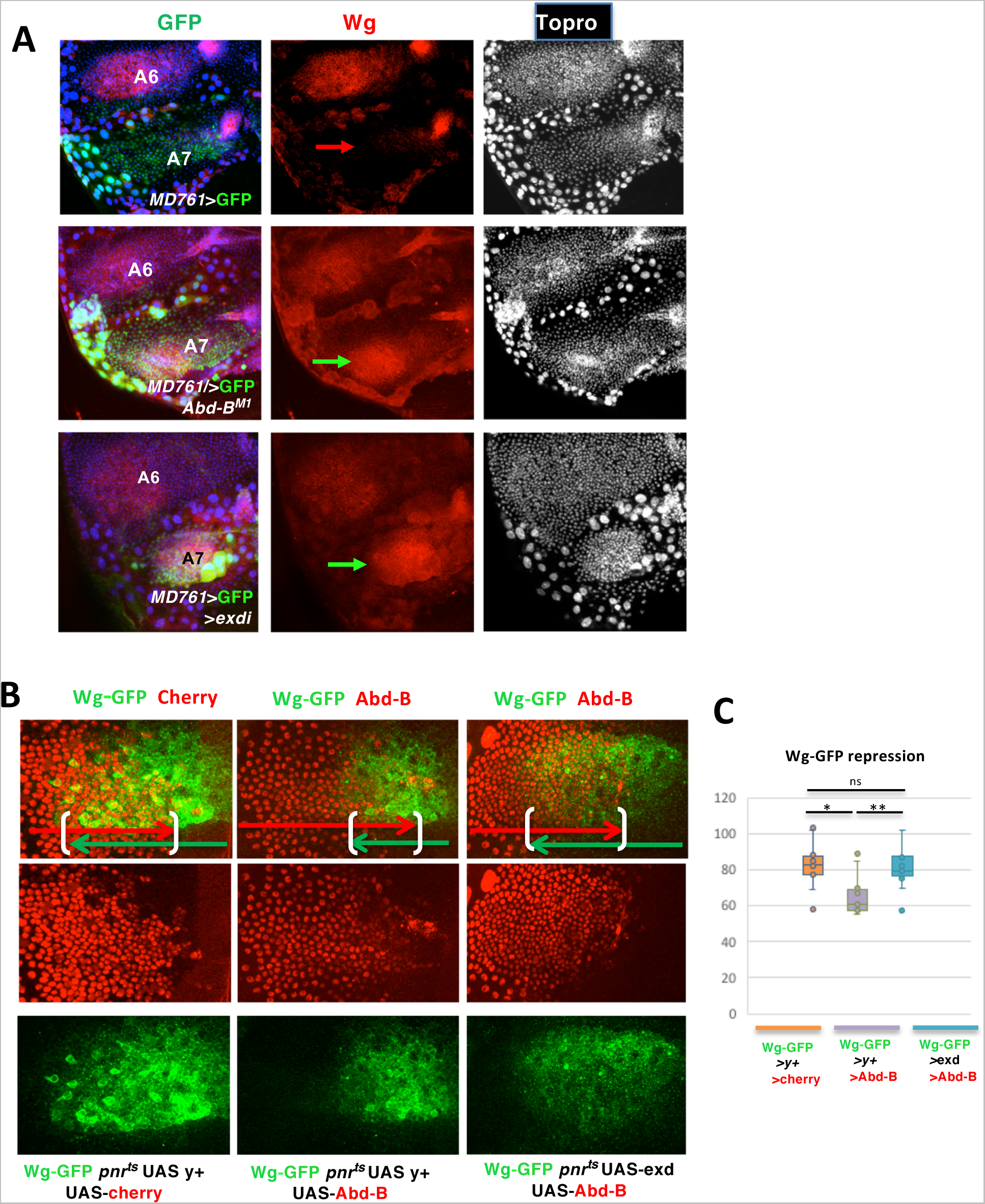
Regulation of wg expression by *exd/hth* and *Abd-B*. **(A)** Expression of Wg in the male A7 of *Abd-B^MD761^* UAS-*GFP*, *Abd-B^MD761^*/*Abd-B^M1^* and *Abd-B^MD761^* UAS-exdRNAi males, showing no Wg expression in the A7 in the first genotype (red arrow) and ectopic Wg expression in this segment of the two latter genotypes (green arrows). **(B)** Expression of wg-GFP in the A4 segment of males of the following genotypes: wg-GFP/UAS-*y^+^*; *pnr*-Gal4 *tub*-Gal80^ts^/UAS-*cherry* (n=10), wg-GFP/UAS-*Abd-B pnr*-Gal4 *tub*-Gal80^ts^/UAS-*y^+^* (n=10) and UAS-*exd*; wg-GFP/UAS-*Abd-B*) *pnr*-Gal4 *tub*-Gal80^ts^/+ (n=12), all shifted from 18°C to 29°C in third larval stage, showing large coincident expression of GFP and cherry in the control, partial repression of wg-GFP when *Abd-B* is expressed in the *pnr* domain and partial suppression of this repression if *exd* is also expressed in this mutant combination. **(C)** Quantification of the extent of overlap of the wg-GFP and cherry, or wg-GFP and Abd-B signals in the three genotypes. Statistical analysis was done by One-way ANOVA.

Collectively, these data set indicate that Abd-B and Exd/Hth function converge in controlling A7 development, including A7 suppression and posterior identity specification, and that accurate levels of both components are required for proper Abd-B activity and A7 development.

### Abd-B associates with Exd/Hth in the pupal abdomen independently of the generic HX interaction motifs

Excess of *exd* or *hth* results in partial loss of A7 suppression, producing an *Abd-B* mutant-like phenotype without lowering *Abd-B* expression levels. This suggests that Exd/Hth reduces Abd-B activity, which could be possibly mediated through physical inhibitory interaction between Exd/Hth and Abd-B proteins, as previously observed (Sambrani et al., 2013; Rivas et al., 2013)

We first probed the possibility of physical interactions between Abd-B and Exd/hth by co-immunoprecipitation after producing the proteins in *Spodoptera frugiperda cells* (*SF*9 baculovirus expression system; Fraser, 1989). We infected SF9 cells with baculovirus expressing tagged Exd, Hth or Abd-B proteins (His::Exd, Flag::Hth and HA::Abd-B) and incubated SF9 extracts on a resin coupled to anti-Flag. After elution we observed co-immunoprecipitation of Abd-B and Exd (Fig. 5A), whereas if Flag::Hth was omitted, neither Abd-B nor Exd was bound by the resin. The amount of Abd-B interacting with Hth in the absence of Exd is much smaller than in its presence. These results indicate that Abd-B associates with Exd/Hth in a predominantly trimeric complex.

**Figure 5.**
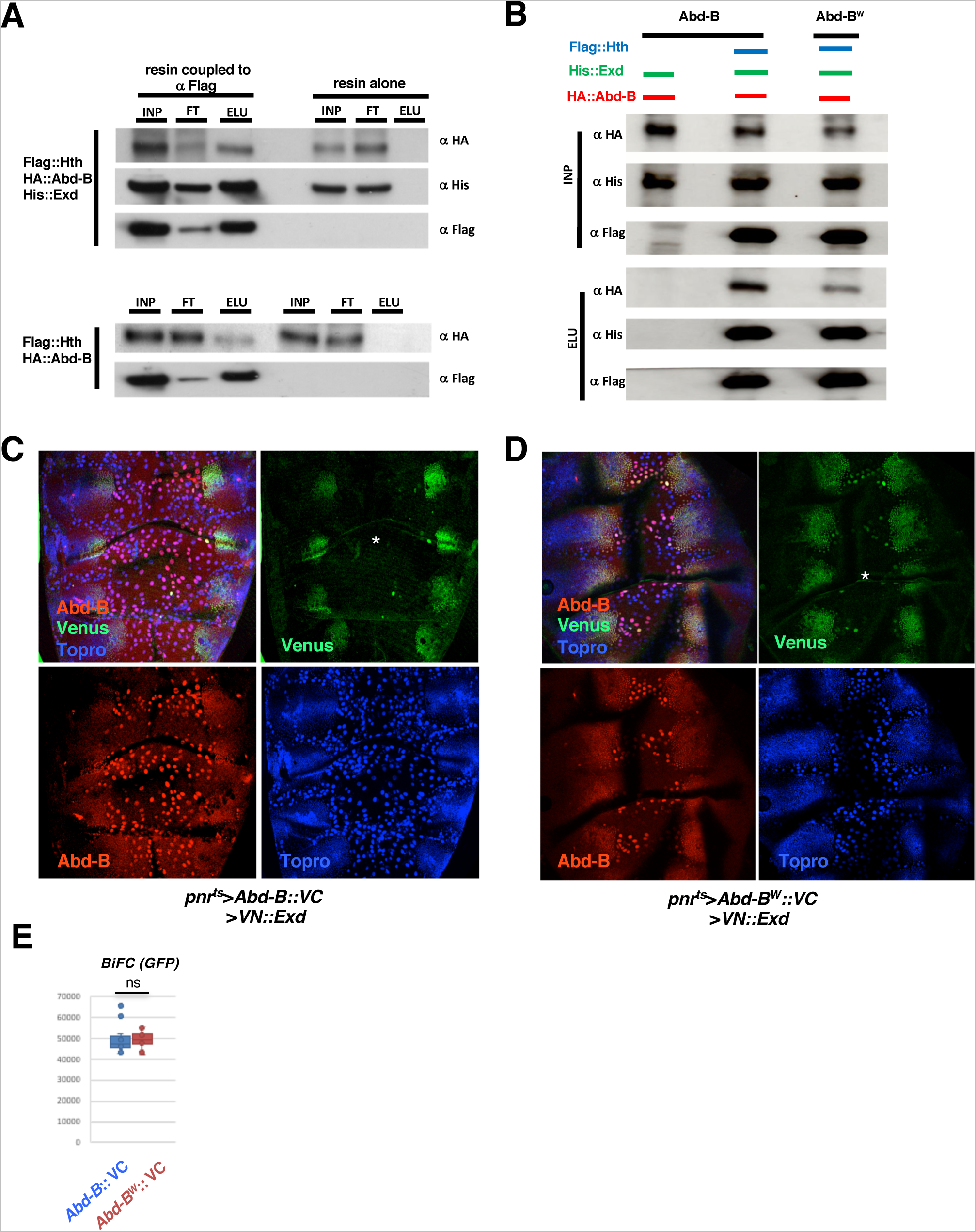
Co-immunoprecipitation and BiFC experiments related to Abd-B and Exd interaction. **(A)** AbdB, Exd and Hth co-immunoprecipitations (co-IPs). INP, FT, and Elu stands respectively for input (the material loaded on the resin), flow through (the material not captured by the resin) and the elution (the material bound to the resin and boiled eluted. Protein present in the INP, FT and ELU are detected by the fused tag: Flag for Hth that also serves for capturing on the anti flag resin, HA for Abd-B and His for Exd. The resin not coupled to anti Flag serves as a control (three right columns). Above is the co-IP in the presence of the Hth, Abd-B and Exd, and below the co-IP in the presence of Hth and Abd-B. Abd-B co precipitates with Hth in the presence and absence (to a lesser extent) of Exd. **(B)** Comparative Abd-B and AbdB ^W^ co-IPs. Protein present in the co-IP are indicated above the gels. There is a significant decrease in the amount of the Abd-B^W^ protein captured on the resin when compared to the Abd-B wildtype protein. **(C)** Dorsal view of a pupa of approximately 24-28h APF of the genotype *pnr*-Gal4 *tub*-Gal80^ts^ UAS-Abd-B::VC UAS-VN::Exd showing Venus signal (in green), Abd-B expression (in red) and Topro, marking nuclei (in blue). See that there is Venus signal, indicating complementation between the Venus fragments of the Abd-B and Exd proteins in histoblasts but not in the polytene LECs (asterisk). **(D)** Dorsal view of an approximately 24-28h APF pupa of genotype *pnr*-Gal4 *tub*-Gal80^ts^ UAS-Abd-B^W^::VC UAS-VN::Exd showing Venus signal (in green), Abd-B expression (in red) and Topro, marking nuclei (in blue). There is Venus signal in just a few of the LECS (the preparation has a reduced number of LECs due to tearing of the tissue during mounting). **(E)** Quantification of BiFC signals. There is similar Venus signal in the experiments performed with the Abd-B and Abd-B^W^ constructs (n=12 for the two genotypes). Statistical analysis was done by two-tailed t-test.

The interaction between Exd and Hox proteins is generally mediated by the Hexapeptide (HX), found in most Hox proteins shortly before the HD (Mann and Chan, 1996). In the Abd-B protein the HX diverges from the canonical core HX sequence, but a Tryptophan that defines the core of the HX motif is present (Shen et al., 1997; Mann et al., 2009). To probe the involvement of the Abd-B HX-like motif in Exd/Hth interactions, we repeated the in vitro co-immunoprecipation experiments using an Abd-B protein bearing a mutation of this Tryptophan towards an Alanine (Abd-B^W^). In these experiments, we observed a large reduction (about 50%) in the amount of co-precipitating Abd-B^W^, compared to wild type Abd-B (Fig. 5B). The Abd-B^W^ protein that remains bound to the Hth-resin could result from Abd-B-Exd interaction not dependent upon the HX like motif, and/or from direct interaction of Abd-B with Hth, as seen in the experiments conducted with the wild type Abd-B protein (Fig. 5A). Together, these data demonstrate that Abd-B associates with Exd/Hth by binding to both proteins, and that the generic HX interaction motif contributes, but is not essential, for these interactions.

We next studied protein interactions directly in the posterior abdomen by Bimolecular fluorescence complementation (BiFC) using the Gal4/UAS system (Hudry et al., 2011). We used UAS constructs in which the coding regions for the N-terminal or C-terminal parts of the Venus fluorescent protein are fused to the coding regions of the Exd or Abd-B proteins (UAS-*VN::Exd* and UAS-VC::*Abd-B*) (Hudry et al., 2011; Hudry et al., 2012). We expressed simultaneously the two constructs in the dorsal abdomen of third instar larvae and pupae with the *pnr*-Gal4 line and the Gal4/Gal80^ts^ system. We observed BiFC in the region of co-expression, the dorsal central abdomen (Fig. 5C), including in the posterior segments where endogenous Abd-B is expressed. Signal resulting from BiFC is almost exclusively confined to histoblasts and not present in LECs, despite expression of the *VN::Exd* and *Abd-B::VC* proteins in both cell types. This result indicates that Abd-B associate with Exd in the pupal abdomen, specifically in histoblasts and not in LECs. To probe for the contribution of the HX-like motif, BiFC experiments were repeated using a *VC::Abd-B^W^* UAS construct inserted at the same chromosomal position, allowing for similar expression levels. Results showed a level of BiFC very similar to that seen with the wild type Abd-B::VC protein (Fig. 5D, E). We concluded that in the posterior abdomen, Abd-B associates with Exd, specifically in histoblasts, and that the core Tryptophan of the HX-like motif does not contribute to this interaction.

### Abd-B protein requirement of A7 suppression: dispensability of the HX central Tryptophan residue but requirement of the HX like motif and posterior-Hox specific protein sequences

While the SF9 co-IP experiments identifies a contribution of the HX-like motif to Exd interaction, the BiFC data in the A7 abdomen indicates that it is not necessary for interaction with Exd, suggesting it may be dispensable for A7 suppression by *Abd-B*. To probe this, we used, as before, a rescuing experiment combining *Abd-B^MD761^*/*Abd-B^M1^*, which alleviates the ability of *Abd-B* to suppress A7 (Fig 3F), with UAS lines allowing for Abd-B or Abd-B^W^ expression (Fig. 6A shows a scheme of the rescuing experiments). In the condition of our experiments (see Methods), the expression of the Abd-B wildtype protein partially corrects the Abd-B mutant phenotype, with an average of 31,6 bristles, while the *Abd-B^MD761^*/*Abd-B^M1^* A7 segments display an average of 65 bristles (Fig. 3F, 6D, E). With the expression levels of wild type and Abd-B^W^ proteins being very similar (Fig. 6C), scoring of A7 bristles indicates that the rescuing ability of A7 suppression by both proteins is also very similar (Fig. 6D, E). This result supports that the Tryptophan residue of the HX-like motif is neither required for Exd interaction in A7 (Fig. 5C), nor for Abd-B A7 suppressive activity.

**Figure 6.**
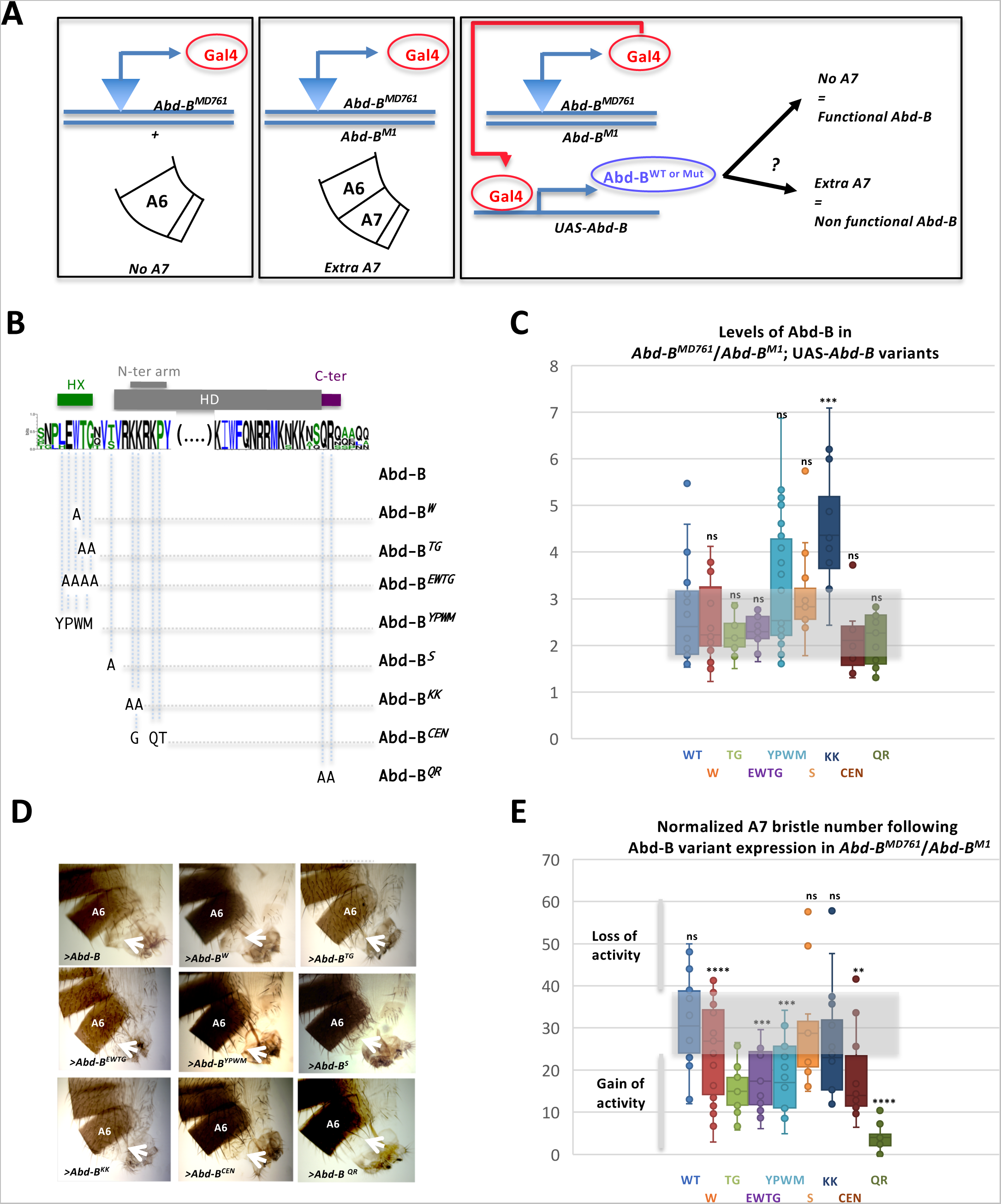
Activity of the wildtype and Abd-B^(W>A)^ mutant proteins. **(A)** Scheme of the genetic experiments performed to ascertain the rescue of the *Abd-B* mutant phenotype of *Abd-B^MD761^* /*Abd-B^M1^* males (A7 transformed into A6) by the expression of wildtype or different *Abd-B* variants. **(B)** Scheme of Abd-B protein mutations. The sequence of the Abd-B protein is indicated, as well as its most conserved domains, the Homeodomain (HD), the Hexapeptide (HX), and sequences immediately C-terminal to the HD. The N-terminal arm of the HD is highlighted. **(C)** Levels of Abd-B proteins expression in the rescue experiment. Levels of the wild type (WT) and different Abd-B variants (referred to according to the labeling in **B**) measured as the amount of protein in a defined area of A7/amount of protein in the same area of A6 dorsal histoblast nests in pupae of about 24-28h APF; n=20 (Abd-B), 19 (W), 11 (TG), 11 (EWTG), 19 (YPWM), 15 (S), 12 (KK), 11 (CEN) and 15 (QR). **(D)** Examples of the posterior abdomens of males of the *Abd-B^MD761^*/*Abd-B^M1^* common genetic background expressing wild type or Abd-B variants (UAS constructs represented as > Abd-B [x]). Arrows point to the A7. **(E)** Quantification (measured as number of bristles) of the rescue of the A7 mutant phenotype observed in *Abd-B^MD761^* /*Abd-B^M1^*animals by the different Abd-B protein variants. Bristle numbers were normalized relative to the level of Abd-B and Abd-B protein variant expression (counts / ratio Abd-B variants/ Abd-B), n=18 (Abd-B), 26 (W), 15 (TG), 16 (EWTG), 21 (YPWM), 11 (S), 16 (KK), 14 (CEN) and 12 (QR) Statistical analysis on normalized bristle number values was performed using the Wilcoxon test.

Facing the lack of requirement of the W residue for Exd-dependent Abd-B function in male A7 suppression, we investigated more broadly possible contributions of other Abd-B protein sequences (Sambrani et al. 2013; Fig. 6B): mutations that target amino acids located within the EWTG HX like motif (W>A, TG>AA, EWTG>AAAA or EWTG>YPWM, the sequence of a canonical HX motif), within the short linker region connecting the HX to the HD (an evolutionarily conserved S>A), within the HD N-terminal arm known to provide paralog specificity (Joshi et al., 2007; Slattery et al., 2011), either mutating the posterior class specific signature (KK>AA) or switching it towards a central class specific signature (CEN: KRKP>CRQT), or within sequences just downstream to the HD (Cter:QR>AA), shown to provide an alternative Exd interaction motif in the central Ubx and Abd-A Hox proteins (Merabet et al. 2007; Saadaoui et al., 2011; Lelli et al., 2011). Sequences mutated are evolutionary conserved in Abd-B proteins from other insects.

Before probing the functional impact of Abd-B mutations, we studied the expression levels of all Abd-B variants in the A7 abdomen (Fig. 6C). Taking the expression level of the *Abd-B^MD761^*-driven wild type protein as a reference point, we found that most proteins are expressed within a similar range of expression level. The most significant exception is the Abd-B^KK^ protein, whose expression is significantly higher than that of the wild type Abd-B protein. Differences in expression levels (subtle as well as stronger differences seen for Abd-B^KK^) were taken into account to normalize the rescuing ability of Abd-B variants (see methods). Results show that none of the mutations results in a weakening of Abd-B activity (Fig. 6D, E). On the contrary, several mutations lead to increased Abd-B A7 suppressive activity. This includes mutations within the HX like motifs (in particular Abd-B^TG^, Abd-B^EWTG^, and Abd-B^YPWM^), within the HD N-terminal arm (Abd-B^CEN^) and in the conserved sequence C-ter to the HD (Abd-B^QR^).

Thus, while the core Tryptophan residue within the HX motif is dispensable for A7 suppression, other residues within the Abd-B HX-like motif are required, as are protein sequences specific to posterior Hox class proteins, in positions known to mediate interaction with Exd in other Hox proteins.

### Context specific usage of intrinsic protein determinants for Abd-B functions

To further probe for a possible function of the W residue of the Abd-B HX like motif in the abdomen, we examined its requirement for development of the female genitalia vaginal teeth, which form two rows easily identifiable (Fig. 7A). Vaginal teeth are influenced by *Abd-B* (reduction of *Abd-B* expression eliminates female genitalia, including vaginal teeth, de Navas et al., 2006; illustrated in Fig. 7B for the *Abd-B^LDN^ /Abd-*B^M1^combination), Hth (*hth*^P2^ mutant clones result in more and disorganized vaginal teeth, Estrada and Sánchez-Herrero, 2001), and Exd (reduction of *exd* produce more, bigger and disorganized vaginal teeth, Fig. 7C).

**Figure 7.**
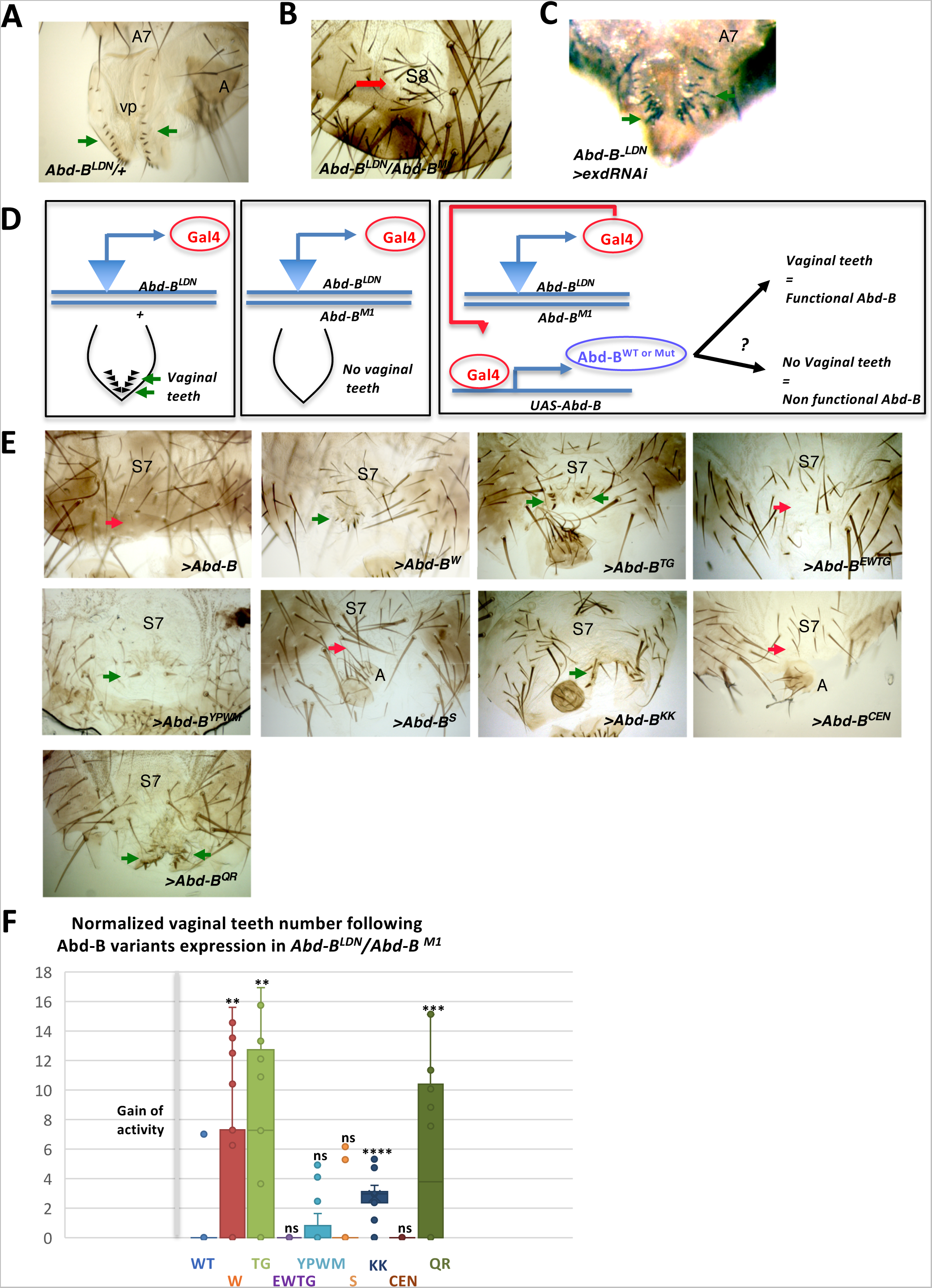
Rescue of the female genitalia mutant phenotype by different Abd-B proteins. **(A)** *Abd-B*^LDN^/+ female genitalia, showing the two rows of vaginal teeth (arrows). **(B)** In *Abd- B*^LDN^/*Abd-B^M1^* females the genitalia is eliminated and frequently replaced by an eighth sternite (S8). **(C)** Reduction of *exd* produces more, bigger and disorganized vaginal teeth**. (D)** Scheme of the genetic experiments performed to ascertain the rescue of the *Abd-B* mutant phenotype of *Abd-B^LDN^* /*Abd-B^M1^* males (elimination of genitalia, including vaginal teeth) by the expression of wildtype or different *Abd-B* variants**. (E)** Examples of female genitalia phenotypes of *Abd-B*^LDN^/*Abd-B^M1^* (common genetic background) expressing wild type or Abd-B variants (UAS constructs represented as > Abd-B [x]) showing no rescue of the mutant phenotype (red arrows) or a small rescue, with presence of some vaginal teeth (green arrows). S7, seventh sternite; A, analia. **(F)** Quantification (measured as number of vaginal teeth) of the rescue of the vaginal teeth phenotype observed in *Abd-B^LDN^* /*Abd-B^M1^* animals by the wild type and Abd-B protein variants; n=24 (Abd-B), 28 (W), 11 (TG), 12 (EWTG), 19 (YPWM), 17 (S), 12 (KK), 13 (CEN) and 20 (QR). Vaginal teeth numbers were normalized relative to the level of Abd-B and Abd-B protein variant expression (counts / ratio Abd-B variants/ Abd-B).

To analyze the function of the different Abd-B variants in female genitalia development we used a rescue experiment similar to the one described for probing restoration of A7 suppression in the male. Instead of the *Abd-B^MD761^* line, we used the *Abd- B^LDN^* line that is mutant for *Abd-B* female genitalia function, and drives expression of Gal4 in female genitalia, mimicking Abd-B expression (see scheme in Fig. 7D). In *Abd-B^LDN^/Abd-B^M1^* females there is a complete lack of vaginal teeth (Fig. 7B), offering the possibility to probe the potential of UAS-driven wild type or mutant Abd-B protein expression to restore vaginal teeth formation. Expression of the wild type Abd-B protein allowed for a very mild rescue, with no teeth in most cases and reaching 7 teeth just in a few individuals (Fig. 7E, F), while the wild type fly harbors around 28 vaginal teeth (Fig. 7A). This poor rescue may be due to this Gal4 line not reproducing precisely in time and space the expression of the wild type *Abd-B*. By comparison, the Abd-B^W^ has a slightly better rescuing ability (Fig. 7E, F). We next investigated the requirement of Abd-B protein motifs already probed in the context of Abd-B A7 suppressive function. As stated above, the wild type protein has a very poor vaginal teeth rescuing ability, only allowing to identify improved activity, which was the major effect seen in the rescue of A7 suppression (Fig. 6). Results identified the TG (within the HX like motif region), KK (within the HD N-terminal arm) and QR (C-Terminal to the HD) mutation as resulting in a major increase of Abd-B vaginal teeth rescuing ability (Fig. 7E, F).

We also probed intrinsic protein requirements for Exd-independent functions. We focused on the wing imaginal disc where Exd is cytoplasmic and therefore inactive (Aspland and White, 1997), investigating in gain of function experiments the expression of *wingless* (*wg*). *wg* is expressed encircling the wing pouch and in the dorso-ventral boundary in the wing disc but not in the posterior compartment of the haltere disc, due to *Ultrabithorax* repression (Weatherbee et al., 1998; Shashidhara et al., 1999). Abd-B protein expression (wild type or mutated version) was driven in the posterior compartment by the *hh*-Gal4 line (Calleja et al., 1996), where its repressive effect on *wg* expression (*wg-GFP* reporter line) was monitored. Results identified a very strong impact of the KK mutations, with the Abd-B^KK^ protein displaying increased *wg*-GFP repressive activity. The TG, YPWM and CEN variants also demonstrate increased *wg*-GFP repressive activity (Supplementary Fig. 4).

The study of intrinsic protein requirements for Abd-B function in the three situations investigated (male A7 suppression, female vaginal teeth development and wing-pouch *wg* repression) shows a clear directionality of observed defects, with mutations in all cases driving increased Abd-B activity. This highlights a prominent role of protein activity buffering, limiting Abd-B activity. We do not know whether these sequences directly mediate contacts towards Exd (and/or Hth). However, the fact that Exd and Hth exert a similar buffering activity suggests that they may do so. If so, and in the case of the HX-like motif, the mode of interaction is likely different as the canonical HX motif, since the central core W residue in Abd-B does not contribute to Abd-B A7 suppressive function while the surrounding residues do. The data also highlight a context specific use of the protein sequences under study: some mutations (TG, KK and QR) impact on the three functions, others (YPWM, CEN) impact two of the three functions and the EWTG mutations specifically impacts on male A7 suppressive function.

### Mutations of the central W residue of the Abd-B like HX motif is required for Abd-B function in the central nervous system

The central W residue of the HX like motif is evolutionary conserved, arguing for functional importance, which contrast with the lack of clear functional requirement We thus aimed at investigating other developmental functions of Abd-B that could reveal a function for the W residue.

Given the contextual use of protein motif described above, we reasoned that the W residue conservation might result from an Exd-dependent function in another tissue than the male abdomen and female genitalia, and examined Abd-B function in the embryonic nervous system. A subset of 30 neuroblasts (NBs) found in each hemi-segment generates the embryonic ventral nerve cord, where segment specific differences are controlled by Hox genes. This is well illustrated by the NB6-4 progenitor, which generates neuronal and glial cells in the thoracic segments, while producing only glial cells in the abdomen. The lack of neuronal cells derived from NB6-4 results from repression by the Hox genes *abd-A* (anterior abdomen) and *Abd-B* (posterior abdomen) (Berger et al., 2005). The abdominal specific NB6- 4 lineage was shown to depend also upon Exd and Hth (Kannan et al., 2010). In the case of *abd-A*, it was further shown that an Abd-A/Exd/Hth complex binds to a *cycE* (*cyclin E*) cis regulatory element mediating transcriptional repression of this gene in NB6-4 (Berger et al., 2005; Kannan et al., 2010). Forcing *abd-A* expression in the thoracic segments was shown to transform the thoracic NB6-4 lineage (neuron + glia) towards an abdominal lineage (only glia) and we have tested if the ectopic expression of *Abd-B* does the same. Wild type or W mutated Abd-B were expressed using the *scabrous* (*sca*)-Gal-4 driver, active from early stages in all neuroblasts, including in thoracic NB6-4. We used Eagle as a marker to identify the NB6-4 lineage (neuron + glia) (Higashijima et al., 1996), Repo as a general marker of glial cells (Xiong et al., 1994; Halter et al., 1995) and the 1.9 *cycE* enhancer shown to recapitulate the Hox-dependent *cycE* expression in NB6-4 (Berger et al., 2005). Results show that *Abd-B* promotes a thoracic to abdominal transformation of NB6-4, evidenced by the lack of NB6- 4/CycE positive cells in the thorax (Berger et al., 2005) (Fig. 8A, B). By contrast, the expression of Abd-B^W^ does not promote such a transformation (Fig. 8A, B), indicating the functional requirement of the W conserved residue of the HX like motif for Abd-B function in controlling segment specificity in the embryonic ventral nerve cord.

**Figure 8.**
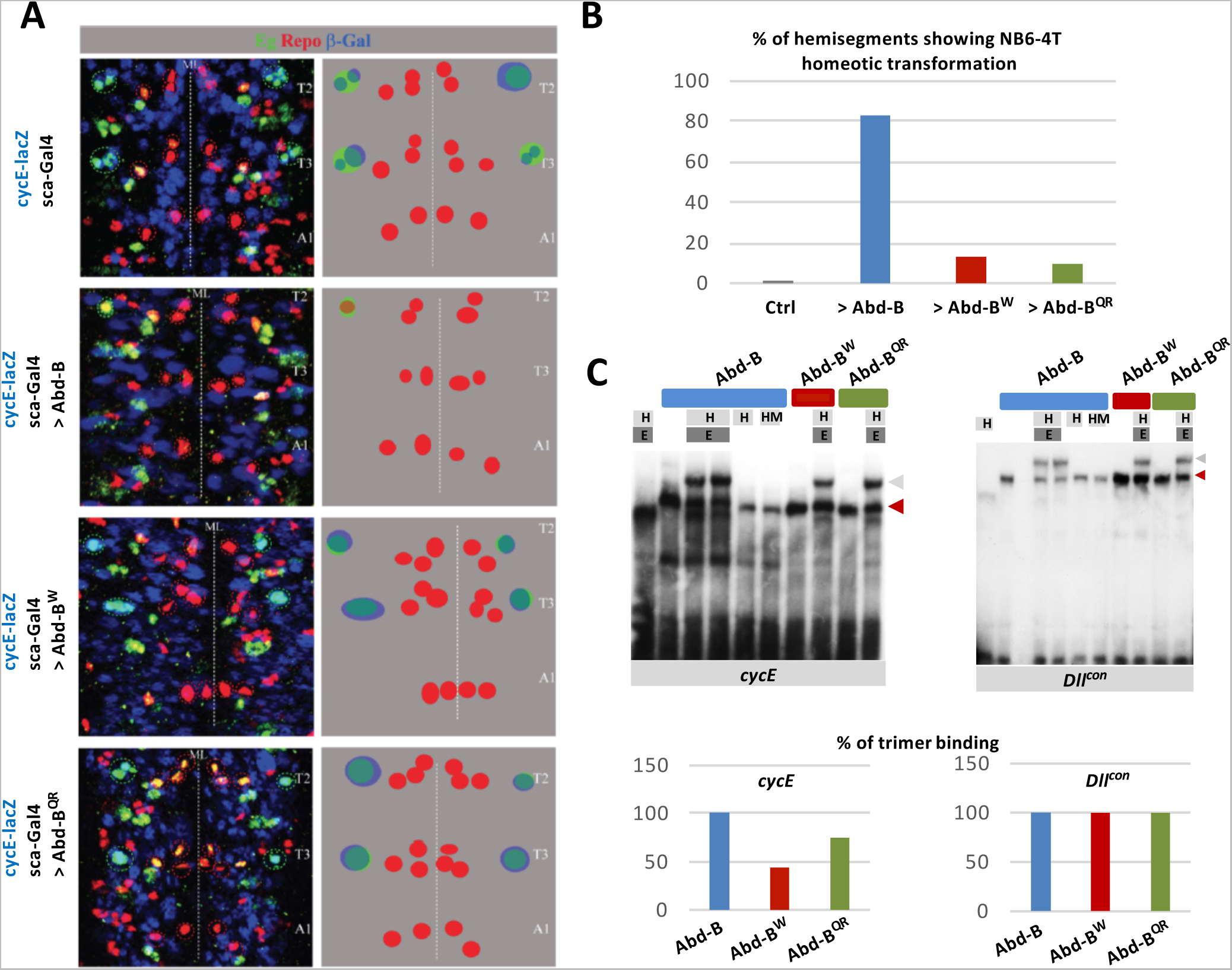
Functional synergy of Abd-B and Exd/Hth in the embryonic CNS. **(A)** Impact of thoracic expression of Abd-B, Abd-B^W^ or Abd-B^QR^ on the NB6-4 CNS lineage. Thoracic expression was achieved using the *scabrous* (*sca*) Gal-4 driver, active from early stages in all neuroblast, including in thoracic NB6-4. Eagle was used as a marker to identify the NB6-4 lineage (neuron + glia) and Repo as a general marker of glial cells. The 1.9 *cycE* enhancer shown to recapitulate the Hox-dependent *cycE* expression in NB6-4 was monitored to assess the impact of Abd-B W and QR mutations on *cycE* expression and NB6-4 abdominal lineage specification. Abd-B promotes a thoracic to abdominal transformation of NB6-4 evidenced by the lack of NB6-4/CycE positive cells in the thorax, a transformation that requires the integrity of the W and QR residues. **(B)** Quantification of the Abd-B, Abd-B^W^ and Abd-B^QR^ abdominal homeotic transformation, assessed by the % of hemisegments showing NB6-4 homeotic transformation. **(C)** EMSA experiments assessing the impact of the W and QR mutation on the formation of an Abd-B/Exd/Hth/DNA complex, using the *cycE* and *Dll^con^* targets. While the W and QR mutations do not affect Abd-B/Exd/Hth/DNA complex formation, they do so on the *cycE* DNA target.

We next investigated if, as shown for Abd-A (Kannan et al, 2010), Abd-B forms a complex with Exd and Hth on DNA elements that mediated Hox repression. Electrophoretic mobility shift assays shows that Abd-B forms a trimeric complex with Exd/Hth on the *cycE* cis-regulatory element (Fig. 8C). Mutation of the W residue affects this trimeric complex with more than half (56%) reduction, while mutation of the QR sequence leads to a 25% reduction (Fig. 8C). Further supporting synergistic DNA binding, we also found that Abd-B and Exd/Hth form a complex on the *DllRcon*, a consensus sequence previously shown to bind most Hox proteins (Gebelein et al., 2002) (Fig. 8C). However, on this specific DNA sequence, the W mutations increases Abd-B monomeric binding, while not affecting the binding of the trimer (Fig. 8C). Taken together with the antagonistic effect Exd/Hth exert on Abd-B binding on DIIR, the sequence mediating repression of the limb promoting gene *Distal-less* (*Dll*; Gebelein et al., 2022; Sambrani et al., 2013), this indicates that both the nature of Abd-B interaction with Exd/Hth and its dependency upon the HX central core W residue depends on the identity of the DNA target sequence. We concluded that in the embryonic ventral nerve cord, Abd-B acts in a manner similar to anterior and central Hox proteins, through synergistic DNA binding with Exd and Hth requiring the HX like motif and specifically the W residue.

We also probed the contribution of the Abd-B QR sequences, located in a position shown to mediate direct contact with Exd in other central and anterior Hox (Foos et al., 2015; Singh et al., 2020). In a manner similar to the W mutation, the QR mutation alleviates Abd-B thoracic to abdominal NB6-4 lineage transformation and impacts the potential of Abd-B to form a trimeric complex specifically on the *cycE* enhancer sequences. This suggest that the mode of cooperative binding used by Abd-B, using both sequences upstream and downstream of the HD, is similar that described for central and anterior Hox proteins.

The posterior Hox genes Ubx, AbdA, and Abd-B were also shown to repress the expression of *eyes absent (eya)*, which is specifically expressed in thoracic neurons. This repression is Exd and Hth dependent (Karlsson et al., 2010). Forced thoracic expression of Abd-B, but not Abd-B^W^, results in the lack of thoracic specific *eya* neurons (Supplementary Fig 5). Similar to the effect on the NB6-4 lineage, Abd-B thus promotes an abdominal transformation that is dependent upon the HX like W central residue. The lack of identified cis regulatory region mediating the Abd-B repression does not allow addressing if this repression also involves the assembling of an Abd-B/Exd/Hth complex on the *eya* cis regulatory region.

## DISCUSSION

In this study we aimed at exploring the nature of posterior class Hox proteins functional interaction with the canonical Pbc/Exd;Meis/Hth cofactors using the *Drosophila* representatives (Abd-B and Exd/Hth) as paradigms and exploring several Abd-B functions. Our data identifies a rich Abd-B cofactor interplay (Fig. 9A) which modulation in different cellular contexts results in a synergistic, antagonistic or dispensable relationship

**Figure 9.**
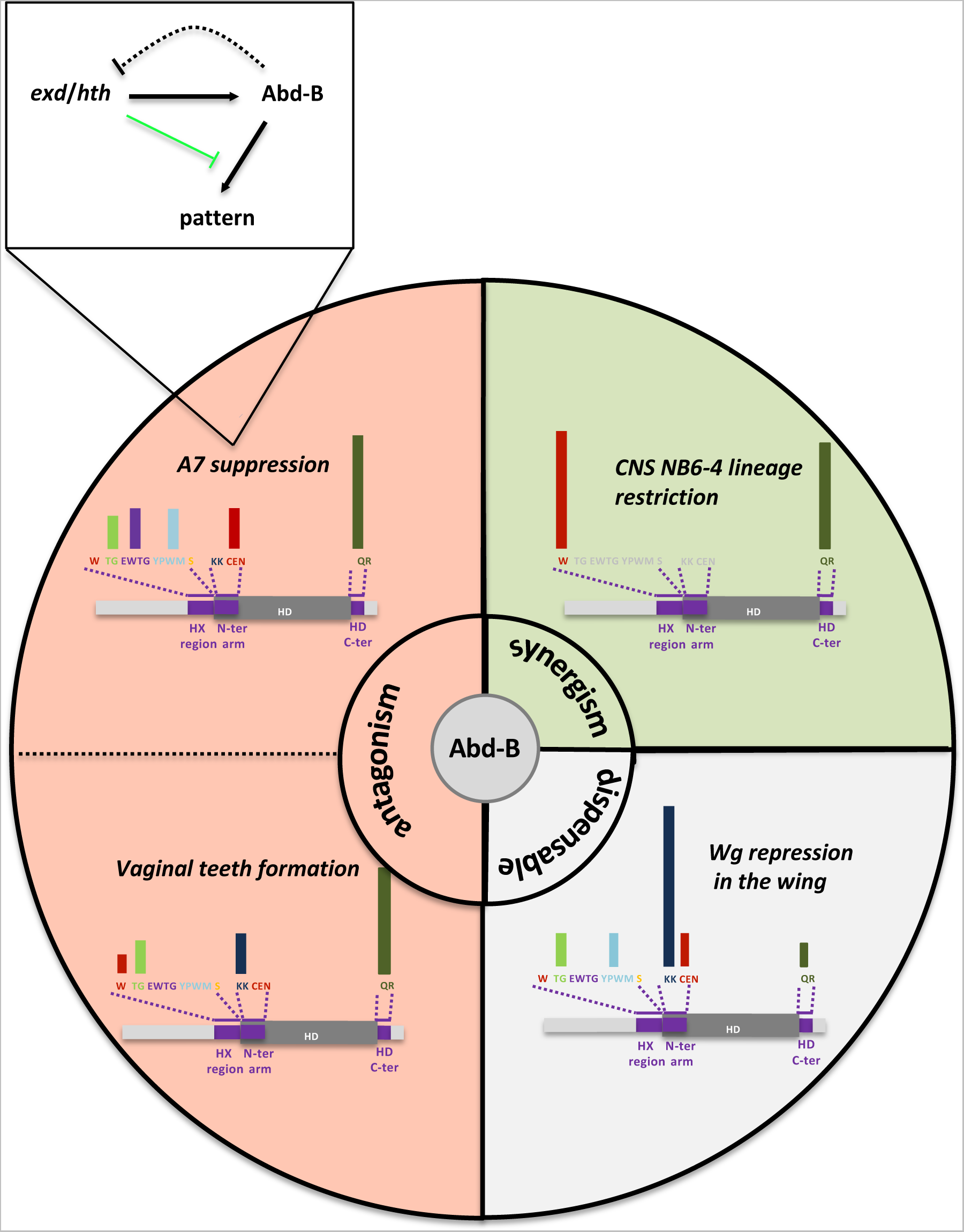
Summary of Abd-B partnership with Exd/Hth and intrinsic protein requirements. Within the circle, functional antagonism is highlighted in red, synergism in green, and dispensability in grey. Protein intrinsic requirement are summarized for each of the four developmental contexts illustrated. Bars above the scheme of Abd- B protein are a qualitative representation of protein sequence requirement. Non-assessed protein sequences are in grey. A functional relationship between Exd/Hth and Abd-B for A7 suppression is indicated.

### Support for Abd-B-Exd/Hth functional antagonism: tight control of Exd/Hth levels prevents Abd-B and Exd/Hth antagonistic action in male A7 suppression

Loss of function experiments showed that loss of *exd* and *hth* mimics *Abd-B* loss in reducing the level of A7 suppression, that is, presenting an A7 segment. Similar results are also observed in the A6 and have been previously described for *exd* (Rauskolb et al., 1995). This primarily suggests a synergistic function of Abd-B and Exd/Hth in A7 suppression, a conclusion also proposed in the case of specification of abdominal denticle fates (Sambrani et al., 2014). However, the reduction of *Abd-B* expression in *hth* mutant clones in the A7 suggests that *exd*/*hth* loss may mimic the *Abd-B* phenotype by impacting on *Abd-B* expression. This is supported by experiments showing the *exd*/*hth* loss of function phenotype is suppressed by increasing *Abd-B* gene dosage. Interestingly, an increase in *exd*/*hth* levels also results in phenotypes resembling *Abd-B* loss, although in this case this is due to an effect on Abd-B activity, not on *Abd-B* expression. In this way, *exd*/*hth* and Abd-B levels are precisely regulated: an increase in *Abd-B* down-regulates *exd*/*hth* expression, and a reduction of *hth* diminishes *Abd-B*, whereas up-regulation of *exd*/*hth* reduces Abd-B activity. The control of Abd-B expression by Exd/Hth, however, does not exclude functional interplay between Abd-B and Exd/Hth at the protein levels, either for synergistic or antagonistic actions.

The positive regulatory interaction between Exd/Hth and Abd-B makes the identification of synergistic action difficult, but allows simpler access to antagonistic actions. Evidence for functional antagonism was obtained from *exd* and *hth* gain of function experiments, that also result in the lack of A7 suppression, but without affecting *Abd-B* expression, indicating that Exd/Hth levels have to be tightly controlled to avoid such antagonistic activity to occur during normal development. Consistently, ectopic *Abd-B* expression down-regulates *wg* expression but this effect is counteracted by an excess of *exd* expression. This highlights the importance of protein dosage, which was also proposed for Ubx protein function, with developmental and evolutionary implications (Paul et al., 2021; Giraud et al., 2021).

Functional antagonism between Abd-B and Exd/Hth was previously described in the embryonic A8 segments, where *Abd-B* promotes the development of the posterior spiracle (Rivas et al., 2013) and represses the limb promoting gene *Dll* (Sambrani et al., 2013). In both instances, *Abd-B* downregulates the expression of *exd*/*hth*, resulting in the lack of Abd- B and Exd/Hth co-existence in the embryonic A8 segment. When Exd and Hth are maintained in this segment they antagonize the activity of Abd-B, impairing posterior spiracle development and *Dll* repression (Rivas et al., 2013; Sambrani et al., 2013). The study of protein binding to a *Dll* regulatory elements that recapitulates *Dll* repression by Abd-B, and to a regulatory element of the *empty spiracle* gene, an Abd-B direct downstream target essential for posterior spiracle development, further indicated that Exd/Hth compete out Abd-B binding to these regulatory elements (Rivas et al., 2013; Sambrani et al., 2013). This was further proposed to result from Abd-B and Exd/Hth direct protein interactions, involving both Abd-B-Exd and Abd-B-Hth interactions, which we also observed *in vivo* in the pupal A7 segments using BiFC, and in SF9 co-precipitation assays. The lack of identified Abd-B direct target sequences mediating Abd-B function in A7 suppression, however, prevents addressing if Exd/Hth counteract Abd-B binding to downstream targets key to A7 elimination. Functional antagonism may also apply in the female genitalia, where Abd-B promotes vaginal teeth formation, while Exd/Hth suppresses them (Estrada and Sánchez-Herrero, 2001).

### Support for Abd-B-Exd/Hth functional synergism: DNA binding cooperativity for NB6-4 lineage restriction

The function of Abd-B in the embryonic CNS provides support for Abd-B-Exd/Hth functional synergism. In this tissue, the NB6-4 progenitor that generates neuronal and glial cells in the thoracic segments will only produce glial cells in abdominal segments. This is due to Ubx/Abd-A function in anterior abdominal segments (A1 to A7), and to Abd-B function in A8 (Berger et al., 2005; Kang et al., 2006). The abdominal specific NB6-4 lineage functionally depends upon Exd and Hth (Kannan et al., 2010). The mechanism of neuronal lineage repression was studied for Abd-A, showing that Abd-A associates with Exd/Hth to bind and repress *cycE*, a key Hox downstream effector for neuronal lineage repression in the embryonic CNS (Kannan et al., 2010). Here we show that, in a manner very similar to Abd-A, Abd-B forms a trimeric complex on a *cycE* regulatory element, indicating that in this specific instance, Abd-B and Exd/Hth cooperative binding mediates functional synergism. Such a functional synergism may also apply to the regulation of the *eya* gene in the embryo, where Abd-B and Exd/Hth are required for repression or gene activation (Berger et al., 2005; Kannan et al., 2010).

Taken together with the above discussed antagonistic function of Abd-B and Exd-Hth, we propose that Abd-B displays both synergistic and antagonistic relationships towards Exd/Hth. The mechanisms channeling Abd-B-Exd/Hth towards synergistic versus antagonistic actions are at least dictated by the identity of the DNA target sequence. This is nicely illustrated by the observation that in in vitro band shift assays Exd-Hth counteract Abd-B binding to a short *Dll* regulatory element (Sambrani et al., 2013), while it cooperatively binds to a slight modification (4 bp) of the same regulatory element (this study). Differences in a cell’s protein content likely may also contribute, which may direct Abd-B and Exd-Hth to different functional outputs in different cellular context, tissues or developmental windows. Although the cooperative or antagonistic effect of Abd-B and Exd/Hth depends on the target, the anteriorwards transformation of A6 and A7 abdominal segments when *exd* or *hth* are up-regulated, including changes in phenotypic traits like pigmentation or formation of trichomes, suggests common antagonistic effects on several genes.

The potentiality of Abd-B to act both synergistically or antagonistically with Exd/Hth is clearly distinct from what the widely supported synergistic functional relationship with anterior and central Hox proteins (Merabet and Mann, 2016). We note, however, that Exd-Hth and the central Hox protein Ubx act antagonistically in *Drosophila* muscle fiber identity specification, with Exd-Hth promoting and the central Hox protein impairing fibrillar (flight muscle) specification (Bryantsev et al., 2012). It was shown that binding of an Exd-Hth complex on *Actin88F (Act88F)*, a key player in fibrillar muscle fate specification, promotes *Act88F* expression and fibrillar muscle identity. In the presence of the central Hox protein Ubx, a trimeric Ubx-Exd-Hth complex assembles on *Act88F*, leading to *Act88F* repression and the establishment of tubular muscle fate as a default state. Although this is a clear case of functional antagonism, it still relies on DNA binding cooperativity highlighting that the output of cooperative DNA binding may be either functional synergism or antagonism.

### Distinct modalities for Abd-B-Exd/Hth interactions for antagonistic versus synergistic functions

The most conserved interaction interface between Hox and Pbc protein lies upstream of the HD, where the HX motif makes major contacts with Pbc class proteins, with a central W docking into a hydrophobic pocket formed by a three amino acid loop insertion specific to Pbc and Meis proteins. Abd-B and vertebrate Hox9-10 (but not Hox11-13 paralogs) have a conserved W just upstream to the HD, but lack canonical HX sequences around this key residue. The study of the contribution of this key W residue for Abd-B and Exd/Hth antagonistic activities in A7 segment suppression largely support dispensability. Further confirming dispensability, in vivo physical interaction between Abd-B and Exd/Hth in A7 segments histoblasts is not impacted by the W mutation. Contrasting with the lack of effects of W mutation, mutations of the HX-like residues TG or EWTG impact A7 suppressive activity, suggesting a mode of Exd/Hth interaction, within the HX region, that may differ from that described for anterior and central Hox proteins. In addition, mutations in the HD N-ter arm, as well as in sequences immediately C-ter to the HD, also impact A7 suppressive activity. Of interest, alternative Pbx interaction modes were identified in the human HoxA9 protein (Dard et al., 2019).

In the case of anterior/central Hox proteins, the Hox/Pbc contacts mediated by the HX influence the positioning of paralog-specific residues lying with the HD N-terminal arm. The KK and CEN mutations that target the Abd-B HD N-terminal arm respectively impact vaginal teeth development and A7 suppressive activity, suggesting a possible concerted action of residues within the HX-like motif and HD N-terminal arm. Of note, Abd-B protein mutations that drive increased protein activity are consistent with Exd/Hth opposing Abd-B function, as altering the interaction through mutation would result in increased activity.

Our work identifies functional synergism between Abd-B and Exd/Hth in the CNS. In both the restriction of the NB6-4 lineage and the repression of *eya*, contrasting with what occurs in the male A7 histoblasts and female genitalia, the W mutation alleviates Abd-B repressive function. In the case of NB6-4 lineage restriction, the regulation of the *cycE* target demonstrates a strict requirement of the W residue for assembling an Abd-B-Exd-Hth complex on *cycE* regulatory region mediating Abd-B repression. This highlights that in synergizing with Exd/Hth, Abd-B seems to use similar molecular modalities than anterior and central Hox protein, with a key contribution of the HX core W residue for interaction with Exd, while it uses different modalities, relying on W surrounding residues, for functional antagonism.

Although less extensively documented, sequences just C-terminal to the HD have been shown to mediate interaction with Pbc class proteins. This was shown for the *Drosophila* central class Hox proteins Ubx and Abd-A (Merabet et al., 2007; Lelli et al., 2011, Foos et al., 2015). Structural and molecular modeling identified a short 8 amino acid region, UbdA, only conserved in Ubx and Abd-A, which establishes highly dynamic contacts with Exd (Merabet et al., 2007; Lelli et al., 2011; Foos et al., 2015). Interestingly, regions C-terminal to the HD are very often highly conserved within Hox paralogous proteins, suggesting that although the molecular modalities of interaction may vary in between paralog groups, the HD C-terminal region may generally be dedicated to contribute additional Pbc contacts. This was recently established for the vertebrate HoxA1 paralog, for which it was further shown that sequence variation in this protein region is key for functional evolution of Hox1 paralog proteins (Singh et al. 2020; Singh and Krumlauf, 2022). The equivalent region in Abd-B is also evolutionary conserved. Mutation of this region in Abd-B (QR) impacts both A7 suppressive function and vaginal teeth development. Here again, the effect of the mutation, resulting in increased Abd-B activity, is consistent with Exd/Hth opposing Abd-B function.

In summary, our data indicate that distinct molecular modalities may underlie Abd-B- Exd/Hth antagonistic and synergistic functions. This is reflected in distinct effects of Abd-B protein mutations for different Abd-B functions, highlighting that a mutation only affects a subset of Abd-B function. This was also observed for the *Drosophila* Abd-A protein, where variable requirements of protein domain for different Abd-A function were reported (Merabet et al., 2011). Additional studies have also shown the effect of mutations in some protein motifs are pleiotropic, with some mutations affecting one character and not others (Hittinger et al., 2005; Prince et al., 2008; Merabet et al., 2011; Sivanantharajah and Percival-Smith, 2009; Sivanantharajah and Percival-Smith, 2014; Sivanantharajah and Percival-Smith, 2015). Differential pleiotropy accounts for the diversity of single Hox proteins and provides a protein landscape prone to evolution. It also highlights that molecular modalities of Hox protein are context specific. The precise understanding of Hox protein mode of action, including how they interface with Pbc/Meis cofactors thus still requires detailed mechanistic analysis of a larger number of Hox protein functions.

## MATERIAL AND METHODS

### Genetics

Vallecas strain was used as a wildtype. *Abd-B^M5^* is a null mutation for the Abd-Bm isoform and *Abd-B^M1^* is a strong loss-of-function *Abd-B* allele for all Abd-B functions (Sánchez-Herrero et al., 1985; Casanova et al., 1986). *hth^P2^* is a strong loss-of-function *hth* allele (Sun et al., 1995; Noro et al., 2006). Wg-GFP is a Wg protein trap (Port et al., 2014) The Gal4/UAS system (Brand and Perrimon, 1993) was used to express ectopically genes or to inactivate them with UAS RNAi constructs. The Gal4/Gal80^ts^ system (McGuire et al., 2003) was used to control the time of activation or inactivation of different genes. In experiments in which larvae or pupae were shifted to 29°C care was taken to take into account the different developmental time at this temperature compared to that at 25°C. We estimated developmental time at 29°C as about 25% longer than at 25°C so that the hours after puparium formation (APF) given are an approximation to the developmental time at 25°C.

#### Gal4 lines

*Abd-B^MD761^* is a Gal4 insertion in the *iab-7* regulatory region of *Abd-B* that drives expression in the A7 segment and is also mutant for the *iab-7* function (named as *MD761*- Gal4 in Foronda et al., 2012). *Abd-B^LDN^* is an insertion in the *Abd-B* gene that reproduces its expression in the female genital primordium (A8) and is also mutant for the *Abd-Bm* function (de Navas et al., 2006; *Abd-B-Gal4^LDN^* in that work). *hh-*Gal4 drives Gal4 expression in the posterior part of the wing disc (Calleja et al., 1996). *sca*-Gal4 (BS 5479) directs expression in all neuroblasts (Mlodzik et al., 1990) and *elav*-Gal4 (elavC155-Gal4, BS 458) in neurons (Luo et al., 1994).

#### UAS lines

UAS-Abd-BM 1.1 (Castelli-Gair et al., 1994), and UAS-Abd-BM (inserted in the ZH35 landing site; Sambrani et al., 2013), UAS*-GFP* (Ito et al., 1997), UAS*-hth* RNAi (Vienna Drosophila RNAi Center, VDRC, 12763, second chromosome and 12764, third chromosome), UAS-*exd* RNAi (VDRC, lines 7802 second chromosome and 7803, third chromosome), UAS*- Abd-B* RNAi (VDRC, line 12024), UAS*-hth* (Pai et al., 1998), UAS*-exd* (Rivas et al., 2013; Sambrani et al., 2013; gift of Natalia Azpiazu, CBMSO, Madrid) UAS-Abd-B^VC^ y UAS-^VN^Exd (Hudry et al., 2012). The different UAS-Abd-B lines were described in Sambrani et al., 2013. Unless otherwise indicated, the UAS-Abd-B line inserted in ZH35 was the one used in the different experiments.

In several experiments we used the Ga4/UAS/Gal80^ts^ system (McGuire et al., 2003) to express different genes during the third larval instar or white pupa stages onwards by shifting the larvae or pupae from 18°C to 29°C. In this way we prevented the lethality, poor viability or abnormal development associated with cell death of different mutant combinations. In the experiments in which we expressed with the *pannier*-Gal4 line different UAS constructs, expressing or inactivating *exd*, *hth* or *Abd-B* (including the BiFC experiments), we shifted the larvae from 18°C to 29°C at the third larval instar and observed pupae normally at about 22-26h APF at 29°C, which we calculated roughly corresponds with about 28-32h a 25°C. In the analysis of rescue of the A7 mutant background (*Abd- B^MD761^*/*Abd-B^M1^*) with the different UAS-Abd-B variants we also used the Gal4/Gal80^ts^ system and shifted white pupae from 18°C to 29°C. The experiments in which we drove expression of the Abd-B variants under the control of the *hh*-gal4 driver in the wing disc, also with *tub*- Gal80^ts^, the shift from 18°C to 29°C was done in second instar or early third instar larvae and fixation of the discs in late third instar larvae. Controls in the different experiments underwent the same temperature treatment.

### Clonal analysis

We used the FLP/FRT system (Golic, 1991; Xu y Rubin, 1993) to make clones mutant for *hth^P2^* or *Abd-B^M5^*. Clones were induced with the hs-flp construct and an hour heat-shock at 37°C during the third instar larval period (although histoblasts do not divide until pupa) and marked by the absence of GFP expression. Flp-out clones (Basler and Struhl, 1993), marked by the presence of GFP expression, were induced during the third instar larval period and induced the expression of *hth, hthRNAi* or *Abd-B*. Flp-out clones were induced by setting the vials at 37°C for 10-15 minute. The chromosomes used to induced clones were: FRT82B *ubi-GFP (*Brand, 1999), *FRT82B hth^P2^* (gift from N. Azpiazu, CBMSO, Madrid). *FRT82B Abd-B^M5^* (Sánchez-Herrero et al., 1985) For flip-out clones, *act>y+>Gal4 UAS GFP* (Ito et al., 1997) was used. The genotypes of the larvae where clones were induced were: *Abd-B* mutant clones: *y hs-flp122/y w o Y; FRT82B Abd-B^M5^/FRT82B ubi-GFP hth* mutant clones : *y hs-flp122/y w o Y; FRT82B hth^P2^ /FRT82B ubi-GFP* flip-out clones: *y hs-flp122/w o Y; act> y+>Gal4 UAS- GFP/UAS-hth RNAi* or *UAS-Abd-B RNAi*.

### Analysis of adult cuticles

Flies were dissected and macerated in a 10% KOH solution at 100°C to remove the internal structures. The cuticles were mounted in glycerol, adding a small amount of Tween-20.

### Inmunohistochemical analysis

Fixations, staining, dissection and mounting of the pupae (male) were done following published protocols (Wang and Yoder, 2011; Foronda et al., 2012). Pupae were visualized in a Zeiss LSM510 and Nikon A1R confocal microscopes. The primary antibodies used were: mouse anti-Abd-B (Hybridoma Bank, Universidad de Iowa), used at 1:10-1:50. Rat anti-Exd (gift of N. Azpiazu, CBMSO, Madrid), (used at 1:100) guinea pig anti-Hth (used at 1:100) (gift of N. Azpiazu, CBMSO, Madrid), rabbit anti-GFP (Invitrogen), (1:300). Secondary antibodies were Alexa 488, Alexa 555, Alexa 647 of the corresponding species (Invitrogen), 1:200. To stain nuclei we used To-Pro 3 (Invitrogen), 1:1000.

Embryo fixing and staining for NB6-4 lineage analysis was as described in Kannan et., 2010.. Primary antibodies used were mouse or rabbit anti-ß-gal (1/1000, Cappel), anti-Repo (mouse, 1/10 DSHB), anti-Eg (rabbit, 1/500, gift of L. S. Shashidhara). The 1.9 kb CycE-lacZ reporter construct strain is defined in Jones et al., 2000. sca-G4, CycE lacZ females were crossed to males of UAS lines. Crosses were maintained all times at 30°C at regular fly culture conditions.

### Bimolecular fluorescence complementation

We have used this method to study interactions between Abd-B and Exd in the dorsal pupal epidermis with the GAL4/UAS system using the *pannier* (*pnr*)-Gal4 line and the Gal4/Gal80^ts^ system, following the methods described before (Hudry et al., 2011), The lines used were UAS-Abd-B^VC^, AS-Abd-B ^(W>A)VC^ and ^VN^UAS-Exd (Hudry et al., 2012; Bischof et al., 2018). Larvae were shifted from 18°C to 29°C at third instar larvae. Quantification of identical areas in the *pnr^+^* and *pnr^-^* domains were done as described above. A minimum of 10 pupae was used to obtain each result.

### Quantification of expression levels

To compare levels of expression of Abd-B of the different Abd-B variants in the experiment of rescue of the A7 segment mutant phenotype we transferred white pupae of the genetic combination UAS-Abd-B (X) *MD761*-Gal4 UAS-GFP *tub*-Gal80^ts^ *Abd-B^M1^* from 18°C to 29°C, let the pupae develop for 24-28h, fixed and stained them with an anti-Abd-B antibody. We selected an area of 10 nuclei (horizontally) per 4 nuclei (vertically) in the posterior part of the A7a and A6a segments, quantified Abd-B levels in both segments with Image J software and obtained the A7/A6 ratio. At least ten pupae were quantified to obtain mean and standard deviation. In this and the rest of quantifications, shifting of larvae or pupae, dissection, fixing and antibody staining was done with the same experimental conditions. Acquisition of the confocal images was also done under identical conditions.

To compare repression of wg-GFP and levels of expression of Abd-B of the different variants when expressed ectopically in the wing disc, we have used the *hh-*Gal4 line (Calleja et al., 1996), which drives expression in the posterior part of the wing disc). We stained the discs with anti-Abd-B and quantified the GFP and Abd-B expression levels with Image J software and the Measure tool. For wg-GFP we selected a small rectangular area encompassing the wg-GFP band of expression at the dorso-ventral boundary in the posterior compartment, adjacent to the A/P compartment boundary, and the same area in the corresponding region of the anterior compartment, also adjacent to the A/P boundary. We measured GFP levels with the Measure tool of image J and divided anterior by posterior levels. A minimum of 10 pupae was used to obtain each result.

In different genetic combinations the levels of Abd-B, Exd or Hth were quantified in pupae also with Image J software. To compare expression levels in the pupal abdomen we used the *pannier* (*pnr*)-Gal4 line, which is driving expression in the central part of the abdomen. The expression domain is labeled with GFP and we have measured, in each case, the expression of Abd-B, Exd or Hth proteins in identical areas in the *pnr^+^* and *pnr^-^*domains (adjacent areas). The measures were taken with Image J software and the relative expression of the *pnr^+^/pnr*^-^ domains is calculated.

### Phenotype normalization

For each variant, the ratio Abd-B variant/Abd-B was determined in the male A7 segment and wing pouch. Given that measurement of protein levels in female genitalia are not possible, we considered that because the exact same UAS-lines were used, the relative expression in the male A7 and female genitalia would be similar (even if absolute levels may differ). The ratio obtained from measurement in the male A7 was this also taken for correcting vaginal teeth counts in the female genitalia. Correction was operated by dividing values per Abd-B variant/Abd-B mean ratio for wg-GFP and vaginal teeth counts, and multiplying values per Abd-B variant/Abd-B mean ratio for A7 bristle counts. Statistical analysis for protein variant activities was performed using the Wilcoxon test (****p<0.0001; 0.0001<***p<0.001; 0.001<**p<0.01; 0.01<*p<0.05).

### Co-immunoprecipitation

The vectors used for the co-immunoprecipitation experiments contain cDNAs encoding the Abd-B (wildtype and different variants), Exd and Hth proteins: fused with hemagglutinin (HA; HA-Abd-B, HA- Abd-B^(W>A)^, Abd-B^(S>A)^ and Abd-B^QRQ>AA)^,) Histidine (His; Exd-His) and Flag (Hth- Flag), respectively. They have been subcloned by conventional procedures in the pFastBac vector (all subclones have been sequenced to verify that they contained the appropriate mutations), and the protocol recommended by Invitrogen (Bac-to-Bac Expression System) has been followed to elaborate baculoviral vectors and induce protein expression. The cells have been collected by centrifugation, washed with cold PBS and resuspended in the lysis solution (140 mM KCl, 5% glycerol, 2.5% MgCl2, 25mM Hepes pH 7.8, Protease inhibitor, 10mM imidazole). Protein expression levels have been quantified by Western Blot, taking these measures into account to mix Exd-Hth with the different Abd-B mutant variants, adding lysis solution where necessary to obtain the same final volume. The immunoprecipitation experiments have been carried out in 2 ml vials, with 200 microliters of anti-Flag resin (Sigma-Aldrich) and 580 microliters of extract. We followed the protocol recommended by the commercial company to carry out the co-immunoprecipitation experiments. The results were visualized by Western Blot, using mouse anti-HA (Sigma-Aldrich), mouse anti-His (Abcam) and rabbit anti-Flag (Sigma-Aldrich and as primary antibodies. Secondary antibodies have been anti-mouse or anti-rabbit coupled to specific IgG light chain HRPs (Jackson). Exposure was made using ECL solution and photographic film.

### EMSA experiments

Abd-B proteins for EMSA were produced using the TNT coupled in vitro transcription/translation system (Promega). Protein production was estimated by labeling the proteins with ^35^S-methionine and found to be produced at similar amounts. EMSAs were performed in 20 µl as described in Pöpperl et al. (1995) using radiolabelled DllR or DllR^con^ (Gebelein et al., 2002) or cycE probes (Kannan et al., 2010).

### Image acquisition

All confocal images were obtained using Zeiss LSM510 and Nikon A1R vertical confocal microscopes. Image treatment and analysis was performed using Image J and Adobe Photoshop and Powerpoint softwares.

### Measurements and statistical analysis

To compare between two groups, as was the case with comparing wildtype Abd-B and the Abd-B^W^ variant (in bristle number, expression levels, BiFC signal), for Abd-B, Hth and Exd levels in different mutant combinations, and for bristle number in comparing flies with different doses of Abd-B and Exd, the two-tailed non-parametric Student’s *t*-test test was used. To compare between more than two groups (wg-GFP repression in pupae expressing combinations of Abd-B and Exd proteins or controls) a non-parametric, one-way ANOVA Dunnett’s test was used. Symbols to indicate significance were : ****p<0.0001; 0.0001<***p<0.001; 0.001<**p<0.01; 0.01<*p<0.05. In all cases the Graph Pad Prism software was used. Raw data for all quantified experiments are displayed in Supplementary Table 1.

## ACKNOWLEDGMENTS

Work in the laboratory of ES was supported by grants from BFU2017-86244-P and PID2020- 113318GB-I00 from FEDER/Ministerio de Ciencia e Innovación-Agencia Estatal de Investigación-Consejo Superior de Investigaciones Científicas, and Work in YG laboratory was supported by the CNRS and AMU, and grants from AMIDEX. We thank L. S. Shashidhara for the anti-Eg antibody, Nuria Prieto for help in some experiments, Flybase, the Bloomington Stock Center, the Vienna Drosophila Resource Center and the Developmental Studies Hybridoma Bank (University of Iowa) for stocks and antibodies.

