## Supplementary material for "Ambivalent partnership of the Drosophila posterior class Hox protein Abdominal-B with the Extradenticle and Homothorax cofactors"

**Supplementary Figures**

**Supplementary Figure 1.** Expression of Exd (**A, A'**) and Hth **(B, B')** in *Abd-B$^{MD761}$* UAS-GFP male pupa of about 28-30h APF, showing similar levels of expression of Exd and Hth in the A7 and A6 segments. *Abd-B$^{MD761}$* is a Gal4 line driving expression in the A7; see Methods and main text).

**Supplementary Figure 2.** Big clone mutant for *Abd-B$^{M5}$* induced in the male A7 segment and marked by the absence of GFP, showing slightly increased levels of Exd (in red) and Hth (in blue) with respect to most adjacent cells.

**Supplementary Figure 3. (A, A')** In *Abd-B$^{MD761}$* UAS-GFP UAS-Abd-BRNAi pupae of about 34-38h APF, in which *Abd-B* expression is reduced in the male A7 segment, the levels of Hth expression are increased with respect to the A6 levels**.** **(B, B')** In *Abd-B$^{MD761}$/+* pupae of about 34-38h APF the levels of expression of Hth in the A7 are also slightly increased as compared to those of the A6.

**Supplementary Figure 4.** Repression of *wg* expression in the wing disc by Abd-B proteins.

**(A-J)** The different panels show wg-GFP expression in *hh*-Gal4 *tub*-Gal80$^{ts}$/+ and in *hh*-Gal4 *tub*-Gal80$^{ts}$ driving expression of the different Abd-B proteins (shifts from 18ºC to 29ºC in second instar larvae). In all cases the larvae were transferred in second or early third instar larvae from 17º to 29ºC. In the wing pouch the wg-GFP line directs GFP expression, like wg, in two rings around the pouch and a dorso-ventral band. (B, C). **(K)** Quantification of Abd-B expression driven by *hh*-Gal4 in the posterior wing pouch. The wild type and Abd-B variant proteins do not show significant differences in expression levels (n=10-15 for the wildtype Abd-B and all the variants). **(L)** Normalized wg-GFP expression following expression of Abd-B and Abd-B variants in the posterior wing pouch. Measurements were taken at the D/V boundary, both in the A (no Abd-B variant expression) and P (Abd-B variant expression) compartments; (n=10-13 for the wildtype Abd-B and all the variants). The A/P ratio of wg-GFP expression is plotted. Significant differences identify gain in Abd-B repressive activity for Abd-$B^{TG}$, Abd-$B^{YPWM}$, Abd-$B^{KK}$, Abd-$B^{CEN}$ and Abd-$B^{QR}$.

**Supplementary Figure 5.** The W residue is required for Abd-B-mediated *eya* repression. Forced *elav*-driven expression of *Abd-B* in the thorax results in the lack of thoracic specific *eya* neurons. Mutation of the W residue alleviates the repression in these neurons.

**Sup. Fig. 1**

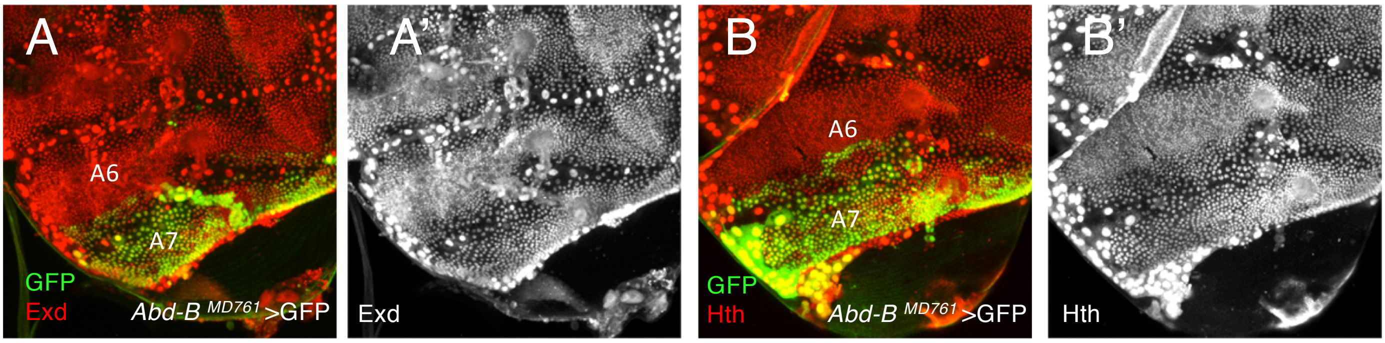
A
A6
A7
GFP
Exd
*Abd-B* $^{MD761}$>GFP
A'
Exd
B
A6
A7
GFP
Hth
*Abd-B* $^{MD761}$>GFP
B'
Hth

### Sup. Fig.2

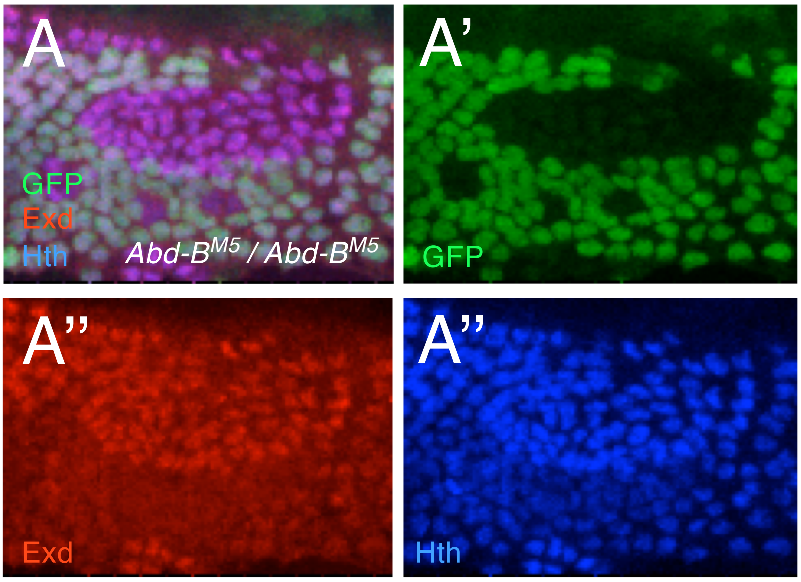

**Sup. Fig.3**

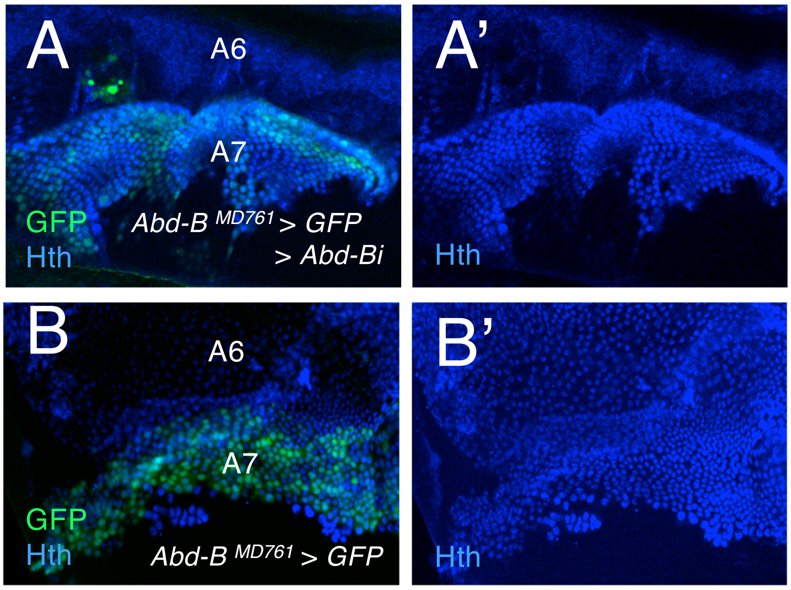
A
A6
A7
GFP
Hth
Abd-B MD761 > GFP
> Abd-Bi
A'
Hth
B
A6
A7
GFP
Hth
Abd-B MD761 > GFP
B'
Hth

#### Sup Fig. 4

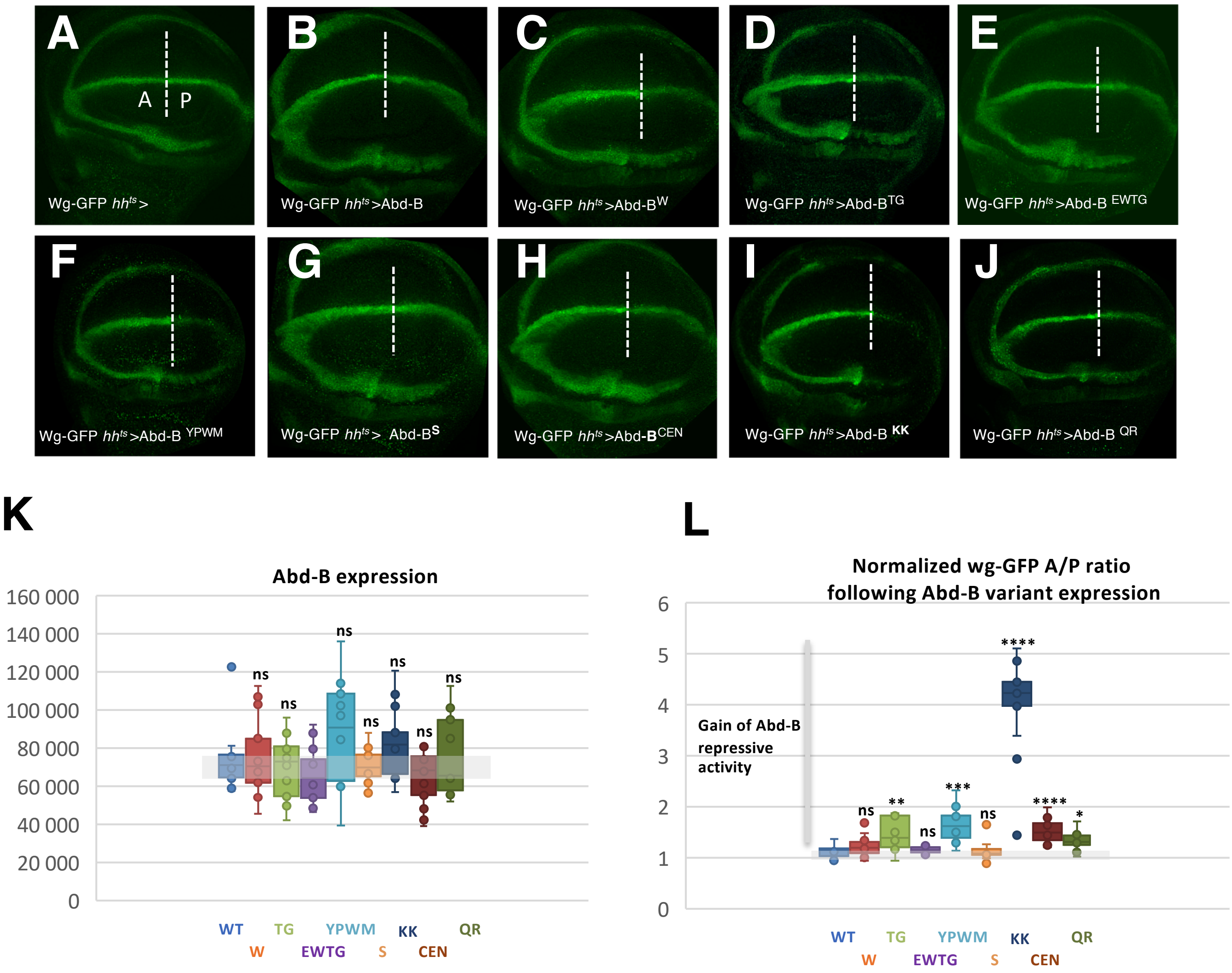
A
A
P
Wg-GFP hh^ts >
B
Wg-GFP hh^ts >Abd-B
C
Wg-GFP hh^ts >Abd-B^W
D
Wg-GFP hh^ts >Abd-B^TG
E
Wg-GFP hh^ts >Abd-B EWTG
F
Wg-GFP hh^ts >Abd-B YPWM
G
Wg-GFP hh^ts > Abd-B^S
H
Wg-GFP hh^ts >Abd-B^CEN
I
Wg-GFP hh^ts >Abd-B KK
J
Wg-GFP hh^ts >Abd-B QR
K
Abd-B expression
160 000
140 000
120 000
100 000
80 000
60 000
40 000
20 000
0
ns
WT
W
TG
EWTG
YPWM
S
KK
CEN
QR
L
Normalized wg-GFP A/P ratio
following Abd-B variant expression
6
5
4
3
2
1
0
Gain of Abd-B
repressive
activity
****
***
**
*
ns
WT
W
TG
EWTG
YPWM
S
KK
CEN
QR

### Sup. Fig.5

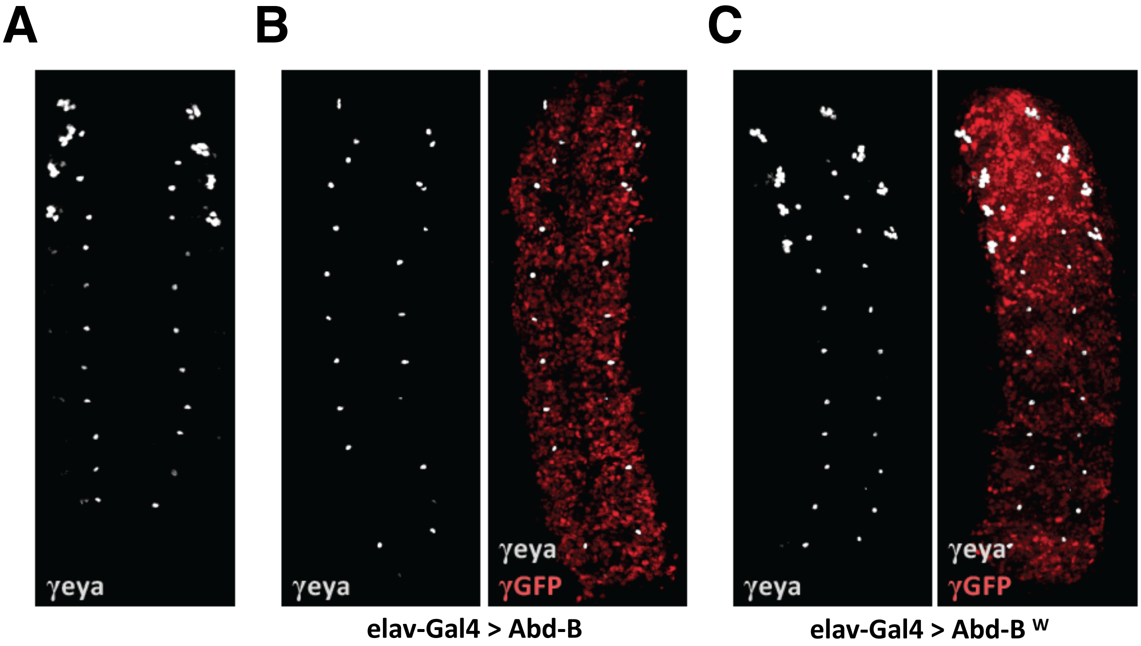
A
B
C
γeya
γeya
γeya
γGFP
γeya
γeya
γGFP
elav-Gal4 > Abd-B
elav-Gal4 > Abd-B W
